# Astrocyte mitochondrial dysfunction and maladaptive lactate supply in multiple sclerosis

**DOI:** 10.64898/2026.11.25.768835

**Authors:** AM Baraibar, T Colomer, N Puente, N Carmona-Molina, A Bernal-Chico, E Sánchez-Martín, I Bonilla-Del Río, C Matute, G Prieto, Y Nasu, E Soria-Gómez, D Jiménez-Blasco, AC Pagano Zottola, I Fernández-Moncada, P Grandes, JP Bolaños, G Marsicano, S Mato

## Abstract

Disruption of metabolic functions underlies the complex responses of astrocytes to neuroinflammation. However, the bioenergetic implications of astrocyte dysfunction in the demyelinating neurodegenerative context of multiple sclerosis remain poorly understood. Here, we investigated the features and mechanistic basis of astrocyte metabolic reprograming in autoimmune demyelinating disease in terms of mitochondrial dysfunction and lactate supply. Ultrastructural analysis revealed that the induction of experimental autoimmune encephalomyelitis (EAE) increased the numbers and reduced the size of mitochondria in cortical astrocytes. Interestingly, we found that astrocytes purified from the brain of EAE mice display significant alterations in the structural organization of their mitochondrial respiratory chain (MRC), as evidenced by decreased proportions of complex I and complex III assembled into respiratory supercomplexes. Consistently, cultured astrocytes activated by proinflammatory cytokines displayed downregulation of genes related to mitochondrial function together with subtle MRC assembly defects and reductions in oxygen consumption rates. Notably, *in vitro* and *ex vivo* imaging of lactate dynamics using genetically encoded biosensors revealed that the astrocytic and extracellular responses to extracellular stimuli mirroring neuronal activity, namely K^+^, NH_4_^+^ and cannabinoids, are impaired by neuroinflammation. At a mechanistic level, reductions in cannabinoid-induced astrocyte lactate flux were associated with downregulated levels of plasma membrane CB_1_ receptors in the brain cortex of EAE mice. Our data reveal mitochondrial structural remodeling and inefficient activity-dependent lactate supply as features of astrocyte bioenergetic reprogramming during autoimmune inflammation and support dysfunction astrocyte-neuron metabolic coupling as pathogenic mechanism of cortical dysfunction in multiple sclerosis.

## Introduction

Among other functions, astrocyte networks sustain brain tissue homeostasis and integrity through the activity-dependent transfer of bioenergetic metabolic messengers (Barros et al., 2025). Astrocytes possess substantial mitochondrial oxidative capacity but, relative to neurons, generally maintain a predominantly glycolytic and metabolically flexible phenotype (Aten et al., 2022; Hertz et al., 2007; Lovatt et al., 2007; Madrer et al., 2025). Indeed, these cells display a unique capability to increase their glycolytic pathways in response to neuronal activity thereby supporting the astrocyte-neuron lactate shuttle model of metabolic coupling (Bonvento & Bolaños, 2021; Bélanger et al., 2011; Magistretti & Allaman, 2018). This distinctive bioenergetic profile relies on the high activity of key glycolytic enzymes as well as in the structural organization of the mitochondrial respiratory chain (MRC) that sustains oxidative phosphorylation (OxPhos) (Almeida et al., 2023). Specifically, astrocytes bear a loosely assembled complex I, whereas neurons assemble complex I to complex III and IV to form respiratory supercomplexes (SCs) that favor electron transport chain efficiency while limiting the production of mitochondrial reactive oxygen species (mROS) (Almeida et al., 2023; Lopez-Fabuel et al., 2016). The complex interplay between mitochondria function and glycolysis that characterizes astrocyte bioenergetics shapes the efficiency of astrocyte-neuron metabolic coupling and its alterations contribute as potential pathogenic mechanisms to numerous neurological diseases (Beard et al., 2021; Bolaños & Magistretti, 2025).

Multiple sclerosis (MS) is a chronic, immune-initiated inflammatory disease of the central nervous system (CNS) and the most common cause of neurological disability in young adults, affecting over 2.8 million people worldwide (Walton et al., 2020). Analysis of MS autopsy material reveals demyelination, glial reactivity and axonal degeneration affecting both white and gray matter of the brain and spinal cord as the main pathological hallmarks of the disease (Lassmann, 2018; Mahad et al., 2015). Both genetic and environmental factors contribute substantially to the complex etiology and heterogeneous clinical presentation of the disease. However, the ultimate causes that drive the development and progression of neurological disability remain elusive. Research conducted over the last years has documented alterations in astrocyte bioenergetic metabolism associated to the acquisition of reactive states as potential pathogenic mechanism in MS (Bittner et al., 2023; Linnerbauer et al., 2020). Early studies targeting mitochondria dysfunction in MS evidenced reductions in mitochondrial gene expression and impaired functional activity of complex I in *postmortem* motor cortex (Dutta et al., 2006). Further analysis suggested the presence of increased mitochondrial numbers in astrocyte profiles within active inflammatory lesions, as compared to normal appearing white matter (Witte et al., 2009). Nonetheless, whether increased astrocyte mitochondrial mass correlates to changes in respiration efficiency and/or in the structural organization of the MRC remains uncertain. Histopathological evaluation of *postmortem* MS tissue also demonstrated upregulated levels of glucose and/or monocarboxylate transporters and glycolytic enzymes in astrocytes within inflammatory lesions (Nijland et al., 2014; Nijland et al., 2015). These anatomical observations suggest an astrocyte glycolytic shift resulting in enhanced astrocyte-axon lactate shuttling that may favor the survival of demyelinated axons. However, functional studies specifically addressing the features and mechanistic basis of astrocyte metabolic adaptations during neuroinflammation offer conflicting observations. *In vitro* approaches suggest that astrocytes reacting to inflammatory stimuli increase their rate of glycolysis, rather than OxPhos, to prevent ATP depletion (Almeida et al., 2004; McComish et al., 2024; Véga et al., 2002). These results fit well with recent transcriptomic data suggesting upregulation of astrocyte glycolytic enzymes in the experimental autoimmune encephalomyelitis (EAE) mouse model of demyelination (das Neves et al., 2021). Conversely, a number of studies in the field report no changes or even deficits in glycolysis and/or lactate flux that are associated to an increased OxPhos performance in astrocytes activated *in vitro* with proinflammatory stimuli (Chao et al., 2019; Gavillet et al., 2008; Meyer et al., 2022). Thus, the metabolic adaptations in inflammatory astrocytes remain controversial. Furthermore, research on astrocyte metabolic contributions to MS focused predominantly on the steady-state bioenergetic consequences of their dysfunction and did not address the dynamics of lactate supply in response to neuronal activity.

Here we addressed the metabolic basis of astrocyte pathology associated to cortical dysfunction in MS. Using biochemical and functional approaches applied to a combination of *in vitro* and *in vivo* models we show that autoimmune inflammation alters astrocyte mitochondrial numbers and morphology, triggers defects in the organization of the astrocyte MRC and impairs lactate efflux in response to physiologically relevant extracellular stimuli that mirror neuronal workload. These findings provide mechanistic insights on astrocyte mitochondrial impairment during neuroinflammation and support a mechanism whereby defects in the neuron-astrocyte metabolic unit may contribute to cortical dysfunction in MS.

## Materials and methods

### Mice

All experiments were performed in accordance with the Guide for the Care and Use of Laboratory Animals (National Research Council Committee, 2011) and the European Communities Council Directive of 22 September 2010 (2010/63/EU74). Animal research procedures were approved by local ethical committees of the University of the Basque Country (approval numbers 2017140, 2017145 and 2022245) and the University of Bordeaux (authorization APAFIS#18111), and the French Ministry of Agriculture and Fisheries (authorization number 3306369). Naive mice on a C57BL/6N background (Janvier, France) and CB_1_^f/f^ mice (carrying the ‘‘floxed’’ *Cnr1* gene) (Marsicano et al., 2002) were used. Cages were enriched and mice were maintained under standard conditions (food and water ad libitum; 12 h-12 h light–dark cycle). Experiments were performed during dark cycle (light off at 8:00 h a.m.). EAE was induced in female mice based on epidemiological evidence that MS affects 2-4 times more women than men (Walton et al., 2020). Primary cultures were prepared from female and male pups obtained from naive pairs. The number of mice in each experimental group was similar. No statistical methods were used to predetermine sample size.

### Drugs

WIN55-212-2 was purchased from Sigma-Aldrich (St-Louis, USA). Tetrodotoxin (TTX), AM251 and AR-C155858 were from Tocris (Bristol, UK). WIN55-212-2, AM251 and AR-C155858 were dissolved in DMSO, TTX was dissolved in ddH_2_O and oxamate was prepared in imaging buffer. The concentration of DMSO was never higher than 0.01%.

### Purified astrocyte cultures

Primary astroglial cell cultures were prepared from 4 to 6-day-old neonatal mouse pups by magnetic activated cell sorting (MACS) as described (Baraibar et al., 2024). Briefly, the forebrains were carefully dissected, meninges removed and tissue enzymatically dissociated using Neural Tissue Dissociation Kit (P) (Miltenyi Biotec). Myelin was removed using a 35% Percoll PLUS (Merck) gradient and cells were sorted using Anti-ACSA-2^+^ MicroBead Kit (Miltenyi Biotec). Cells were seeded onto 6-well plates at a density of 500.000 cells/well for RNA sequencing, and of 1.000.000 cells/well for mitochondrial isolation. Alternatively, astrocytes were seeded onto Seahorse 96-well plates (Agilent) at a density of 50.000 cells/well for metabolic experiments and onto 48-well plates at a density of 50.000 cells/well for extracellular lactate determinations. Cells were maintained in serum-free defined medium containing 50% neurobasal, 50% DMEM, 100 U/mL penicillin, 100 μg/mL streptomycin, 1 mM sodium pyruvate, 292 μg/mL l-glutamine, 1×SATO and 5 μg/mL of N-acetyl cysteine. This medium was supplemented with the astrocyte-required survival factor HBEGF (5 ng/mL; Preprotech). Astrocyte cultures were switched to 10 mM glucose growth medium prepared using 50% neurobasal (Gibco, A2477501) and 50% DMEM (Gibco, 11520416) for 24 h before stimulation. The purity of the cultures was routinely assessed by examining the characteristic cell morphologies under phase-contrast microscopy.

### Mixed cortical glia-neuron cultures

Mixed cortical cultures of neuronal and glial cells were prepared from 1 to 3-day-old neonatal mice (Bittner et al., 2010; Bittner et al., 2011). Briefly, mice were euthanized, the brain removed, and cortex dissected in iced-cold Hank’s balanced salt solution. The tissue was enzymatically digested with trypsin/EDTA for 5 min at 37°C and the enzymatic digestion was stopped with 10% FBS in B-27 supplemented neurobasal medium. After this, a gentle dissociation of the tissue was performed by repeatedly passing it through a 1-mL micropipette tip. Obtained cells were left in suspension to allow debris precipitation and removal. Cells were seeded in 18-mm glass coverslips treated with poly-L-lysine and incubated for 90 min to allow cell adhesion. After this, the medium was replaced with fresh B-27 supplemented neurobasal medium with 10 mM glucose, 0.24 mM pyruvate, 2 mM GlutaMAX^TM^, 100 U/mL penicillin, 100 µg/mL streptomycin and 2.5 µg/mL amphotericin B at 37°C in a humidified atmosphere of 5% CO_2_. At 13-14 days *in vitro* (DIV), cultures were exposed to 1 × 10^6^ plaque-forming units (pfu) of adenoviral vectors (serotype 5) coding for Laconic (San Martín et al., 2013). Measurements were carried out 48-72 h after infection of cells (16-17 DIV). Adenoviral vectors encoding the FRET biosensor were custom-made by Vector Biolabs (PA, USA).

### Cell culture stimulation

Cell cultures were stimulated with 25 ng/mL TNFα (Cell Signaling Technology, 8902SF), 3 ng/mL IL-1α (Sigma, I3901) and 400 ng/mL C1q (MyBioSource, MBS143105) or vehicle for 18-24 h (Baraibar et al., 2024; Liddelow et al., 2017).

### Analysis of extracellular flux

Analyses of mitochondrial respiration and glycolytic rates were performed using the Seahorse XFe96 extracellular flux analyzer (Agilent Technologies) according to the manufacturer’s specifications. The day of the experiment, cells were washed and the medium was exchanged with XF Basal medium supplemented with 10 mM glucose, 2 mM l-glutamine, 1 mM sodium pyruvate (pH 7.4). Cells were then transferred to a CO_2_-free incubator and maintained at 37°C for 1 h before starting the assay. Oxygen consumption rate (OCR) was measured using Seahorse XF Cell Mito Stress Test Kit (Agilent Technologies, 103015-100) following the sequential injection of oligomycin (2 µM) to inhibit ATP synthase, carbonyl cyanide-4 (trifluoromethoxy) phenylhydrazone (FCCP; 1 µM) to induce mitochondrial uncoupling and rotenone/antimycin A (0.5 µM) to inhibit mitochondrial complex I and III, respectively. Glycolysis-derived proton efflux rate (GlycoPER) was estimated using the XF Glycolytic Rate Assay Kit (Agilent Technologies, 103344-100) after the sequential treatment with rotenone/antimycin A (0.5 μM) and the glucose analogue 2-deoxyglucose (50 mM). At the end of the assays, the medium was carefully aspirated and cells incubated with Calcein-AM (Invitrogen, C3100MP) to adjust for potential differences in cell density.

### Lactate release measurements

Following stimulation with proinflammatory cytokines or vehicle, cells were washed and treated for 1 h with oligomycin or vehicle in 10 mM glucose growth medium. Media were collected, centrifuged at 500 x *g* for 5 min at 4°C and stored at −80°C until further use. Extracellular lactate was quantified using the lactate assay kit (Sigma-Aldrich, MAK064) and a Synergy-HT microplate reader (Biotek) according to the manufacturer’s protocol. The absorption signal was normalized to total protein content estimated with the BioRad Protein Assay Kit (Protein Assay Reagents, 5000-114-13-15) using BSA as standard.

### RNA sequencing

#### RNA sampling and sequencing

Total RNA from purified astrocyte cultures was extracted using RNeasy Plus Micro kits (Qiagen, 74034) according to the manufacturer’s instructions and stored at −80°C until analysis. RNA samples were sent for sequencing to the Genomic Platform of the CIC BioGUNE (Derio, Spain). RNA quality and integrity were evaluated with an Agilent RNA 6000 Pico Chips (Agilent technologies, 5067-1513) and the Agilent 2100 Bioanalyzer (Agilent technologies). Sequencing libraries were constructed using 6-8 ng total RNA per sample with the SMARTerStranded Total RNA-seq Kit v3 - Pico Input Mammalian (TakaraBio, 634487) and the SMARTer RNA Unique Dual Index Kit-24U (TakaraBio, 634451). Amplified libraries were visualized on an Agilent 2100 Bioanalyzer using Agilent High Sensitivity DNA Chip (AgilentTechnologies, 5067-4626) and quantified using the Qubit^®^ dsDNA HS assay kit (ThermoFisherScientific, Q32854). Sequencing was performed in the NovaSeq 6000 (Illumina) in 100 bp paired-end reads per sample.

#### Differential gene expression and gene set enrichment analysis

Differential gene expression analysis was conducted on raw count data using the DESeq2 package in R (Love et al., 2014). Log fold changes were shrunk using the approximate posterior estimation for generalized linear model (*apeglm*) (Zhu et al., 2019) method to reduce variance for genes with limited information for statistical inference. ENSEMBL gene identifiers were annotated to gene symbols with the BiomaRt package (Durinck et al., 2009), querying the *Mus musculus*_gene_ensembl dataset from Ensembl version 102. Results were considered statistically significant for *P* values < 0.05 and the absolute value of the log_2_ fold-change ≥ 2.0.

Gene set enrichment analysis (GSEA) (Subramanian et al., 2005) was carried out with gene sets derived from the molecular signatures database (MSigDb) (Hallmarks) (Liberzon et al., 2015) and genome-scale metabolic models (GEM) (Robinson et al., 2020; Wang et al., 2021) using gene set permutations (*n* = 1000) for the assessment of significance and signal-to-noise metric for ranking genes.

### Live cell fluorescence imaging

Mixed cortical glia-neuron cultures were mounted in an open chamber and imaged on wide-field mode with an inverted Leica DMI 6000 microscope (Leica Microsystems, Wetzlar, Germany) equipped with a resolutive HQ2 camera (Photometrics, Tucson, USA). The illumination system used was a Lumencor spectra 7 (Lumencor, Beaverton, USA). The objective used was HCX PL APO CS 40× oil 1.25 NA. Multi-positions were done with a motorized stage Scan IM (Märzhäuser, Wetzlar, Germany). A 37°C atmosphere was created with an incubator box and an air heating system (Life Imaging Services, Basel, Switzerland). The system was controlled by MetaMorph software (Molecular Devices, Sunnyvale, USA). Cells were superfused with an imaging solution consisting of (in mM): 10 HEPES, 112 NaCl, 24 NaHCO_3_, 3 KCl, 1.25 MgCl_2_, 1.25 CaCl_2_, 2 glucose, 0.5 sodium lactate and bubbled with air/5% CO_2_ at 37°C, at a constant flow of 3 mL/min. Astrocytes expressing Laconic were imaged at 40X, and excited at 430 nm for 0.01-0.05 s, emissions collected at 465-485 nm for mTFP and 542-556 nm for Venus, with image acquisition every 10 s. Cultures were sequentially exposed to a rise in extracellular K^+^ (6 mM) and NH_4_^+^ (5 mM) as intercellular signals that trigger astrocyte lactate release (Barros et al., 2023; Lerchundi et al., 2015; Sotelo-Hitschfeld et al., 2015). Cells were subsequently perfused with sodium oxamate (6 mM - in absence of glucose) to halt lactate production followed by the monocarboxylate transporters (MCTs) blocker AR-C155858 (1 μM) in the presence of high lactate concentrations (10 mM) to boost intracellular accumulation (Lerchundi et al., 2015). Alternatively, cultures were exposed to the cannabinoid agonist WIN55,212-2 (1 μM) to stimulate lactate production by CB_1_ receptors (Fernández-Moncada et al., 2024) followed oxamate and AR-C155858 as described. The ratio between mTFP and Venus was computed and normalized to the baseline. For quantification, the mTFP/Venus ratio was transformed to lactate concentration, as described previously (San Martín et al., 2013)

### EAE model

Mice were immunized in the flank by subcutaneous (s.c.) injection of 200 μg MOG_35-55_ peptide (MEVGWYRSPFSRVVHLYRNGK) (Peptide Synthesis Core Facilities of the Pompeu Fabra University, Spain) in incomplete Freund’s adjuvant supplemented with 8 mg/mL Mycobacterium tuberculosis H37Ra (Difco Laboratories). Pertussis toxin (500 ng; Calbiochem) was injected i.p. on the day of immunization and again at 2 days post-immunization (dpi). Body weight and motor symptoms were recorded daily and scored from 0 to 8 as follows: 0, no detectable changes in muscle tone and motor behavior; 1, flaccid tail; 2, paralyzed tail; 3, impairment or loss of muscle tone in hindlimbs; 4, hindlimb hemiparalysis; 5, complete hindlimb paralysis; 6, complete hindlimb paralysis and loss of muscle tone in forelimbs; 7, tetraplegia; and 8, moribund.

### Immunomagnetic purification of astrocytes from adult mice

Mouse adult forebrain tissue was digested in Earle’s balanced salt solution (EBSS) containing 116 mM NaCl, 5.4 mM KCl, 1.5 mM MgSO_4_, 26 mM NaHCO_3_, 1.01 mM NaH_2_PO_4_·2H_2_O, 4 mM glucose, 10 mg/L phenol red (pH 7.2) supplemented with 14.4 μl/mL albumin and 26 μl/mL DNase type I and 10.8 μl/mL trypsin. Dissociated cells, after removal of debris and red blood cells, were separated using an astrocyte-specific anti-ACSA-2 Microbead Kit according to the manufacturer’s protocol (Miltenyi Biotec, 130-097-679). Cells pellets were stored at −80°C until further use.

### Surgery for AAV administration

Mice were anaesthetized with isoflurane and placed on a heating-pad to keep the body temperature at 37°C. Eye dehydration was prevented by topical application of ophthalmic gel and analgesia was achieved by s.c. injection of buprenorphine (Buprecare, 0.05 mg/Kg). The skin above the skull was shaved with a razor and disinfected with modified ethanol 70% and betadine before an incision was made. Mice were placed into a stereotaxic apparatus (David Kopf Instruments) with mouse adaptor and lateral ear bars and injected with AAV5-gfaABC1D-eLACCO3M or AAV5-gfaABC1D-deLACCO3ctrol (1.2 x 10^13^ GC/mL) alone or in combination with ssAAV9/2-hGFAP-mCherry_iCre-WPRE-hGHp(A) (ETH Zurich; 5.2 x 10^12^ GC/mL) to carry out two-photon imaging of extracellular lactate in astrocytes. Stereotaxic injections (600 nL) were targeted to the mouse somatosensory cortex according to the following coordinates (from bregma): anterior-posterior −1.5; medial-lateral ± 2.5; dorsal-ventral −1.3. Viral particles were injected at a maximum rate of 1 nL/sec using a glass pipette attached to a Nanojet III (Drummond, Broomall, USA). Following virus delivery, the syringe was left in place for 10 min before being slowly withdrawn from the brain. Mice were weighed daily and individuals that failed to return to their pre-surgery body weight were excluded from subsequent experiments.

### *Ex vivo* lactate imaging

#### Cortical slice preparation

Mice were euthanized by decapitation and brains were rapidly removed and placed in ice-cold artificial cerebrospinal fluid (aCSF). Coronal brain slices (350 µm-thick) containing the somatosensory cortex were prepared using a Leica VT1200 vibratome in an ice-cold high-sucrose solution containing (in mM): 189 sucrose, 10 glucose, 26 NaHCO_3_, 3 KCl, 5 MgSO_4_, 0.1 CaCl_2_, and 1.25 NaH_2_PO_4_, continuously bubbled with 95% O_2_/5% CO_2_. Following cutting, slices were allowed to recover at room temperature in aCSF containing (in mM): NaCl 124, KCl 2.69, KH_2_PO_4_ 1.25, MgSO_4_ 2, NaHCO_3_ 26, CaCl_2_ 2 and glucose 10, gassed with 95% O_2_/5% CO_2_ (pH = 7.3-7.4). For imaging experiments, slices were transferred to an immersion recording chamber and superfused at room temperature with oxygenated aCSF at a flow rate of 0.5 mL/min.

#### Two-photon fluorescence imaging

Two-photon microscopy imaging of layer V-VI astrocytes in the somatosensory barrel field was performed using a Femto-2D microscope equipped with a tunable Ti:Sapphire MaiTai DeepSee laser (Spectra Physics) and GaAsP PMT detectors. The laser was tuned at 920 nm with a 20X water immersion lens (1.00 N.A.; Olympus) and a 490/60 nm filter. Videos were obtained at 512 × 512 resolution with a sampling interval of 1 s. Imaging experiments were performed in the presence of TTX (1 µM) by bath application of extracellular solutions containing high K^+^ (12 mM), NH_4_^+^ (6 mM) and WIN55,212-2 (3 µM) alone or in the presence of the CB_1_ antagonist AM251 (2 µM). In the high K⁺ solution NaCl was reduced accordingly to maintain osmolarity. Analysis and quantification of fluorescence levels in astrocytes was performed with a custom MATLAB program (Calsee: https://www.araquelab.com/code/). Fluorescence variations recorded at the soma and processes of the cells (ΔF/F_0_) were estimated as changes of the signal during the final 30 s of a 5 min stimulation over a 3 min baseline period.

### Mitochondrial measurements

#### Mitochondria isolation and solubilization

Mitochondria were obtained from astrocyte cell cultures, brain cortical tissue or astrocytes purified from mouse forebrains, according to previously published protocols (Jimenez-Blasco et al., 2020; Vicente-Gutierrez et al., 2019). Briefly, cell pellets and tissue samples were collected, frozen at −80°C and homogenized (15 strokes) in a glass-Teflon Potter homogenizer in buffer A (83 mM sucrose and 10 mM MOPS; pH 7.2). The same volume of buffer B (250 mM sucrose and 30 mM MOPS) was added to the sample, and the homogenate was centrifuged (1,000 x *g* for 5 min) to remove unbroken cells and nuclei. Centrifugation of the supernatant was then performed (12,000 *x* g for 3 min) to obtain the mitochondrial fraction, which was washed in buffer C (320 mM sucrose, 1 mM EDTA and 10 mM Tris-HCl; pH 7.4). Finally, mitochondria were suspended in buffer D (1 M 6-aminohexanoic acid and 50 mM Bis-Tris-HCl; pH 7.0) and quantified using the Pierce™ BCA Protein Assay Kit (ThermoFisher, A55864).

#### Blue native gel electrophoresis (BNGE)

To assess the mitochondrial respiratory complexes, digitonin (Sigma-Aldrich D141) (4 g/g)-solubilized mitochondria (10-50 μg) were loaded in NativePAGE Novex 3-12% gels (Life Technologies, 10024112). After electrophoresis, in-gel NADH dehydrogenase activity (IGA) was evaluated and the individual complex I and complex I-containing SC bands were identified due to the formation of purple precipitates (Lopez-Fabuel et al., 2016). Briefly, gels were incubated in 0.1 M Tris-HCl buffer (pH 7.4), 1 mg/ml of nitro blue tetrazolium (Sigma-Aldrich, N5514) and 0.14 mM ß-Nicotinamide adenine dinucleotide reduced disodium salt hydrate (NADH) (Sigma-Aldrich, N8129). Direct transfer was also performed after soaking the gels for 10 minutes in methanol followed by 10 minutes in carbonate buffer (10 mM NaHCO_3_, 3 mM Na_2_CO_3_·10 H_2_O; pH 9.5-10). Protein transfer to PVDF membranes (Thermo Scientific, 88520) was carried out at 300 mA and 60 V for 1.5 h at 4°C in carbonate buffer. Membranes were blocked for 1 h in Tris-buffered saline (TBS) (50 mM Tris, 200 mM NaCl; pH 7.4) containing 0.05% Tween-20 (TBS-T) with 5% BSA and incubated overnight at 4°C with primary antibodies (**Table 3**). After 3 washes in TBS-T membranes were incubated with horseradish peroxidase-conjugated secondary antibodies (1:5000; Cell Signaling Technology) for 1 h at RT and developed with NZY Standard (Nzytech, MB40101) or NZY Advanced (Nzytech, MB40201) ECL Western Blotting Substrates. Analysis of relevant immunoreactive bands was carried out using Fiji Image J software after acquisition on a ChemiDoc XRS System (Bio-Rad). Protein bands from complex II SDHA subunit were used as loading controls for signal normalization.

### Electron microscopy

Mice were deeply anesthetized by intraperitoneal (i.p.) injection of ketamine/xylazine (80/10 mg/Kg body weight) and were transcardially perfused at room temperature (RT; 20-25°C) with 0.1 M phosphate buffered saline (PBS; pH 7.4) for 20 s, followed by a fixative containing 4% formaldehyde (freshly depolymerized from paraformaldehyde), 0.2% picric acid, and 0.1% glutaraldehyde in 0.1 M phosphate buffer (PB; pH 7.4) for 10-15 min. Then, brains were removed from the skull and post-fixed in the same fixative for about 1 week at 4°C. Afterwards, brains were stored at 4°C in 1:10 diluted fixative solution until used.

Fifty µm-thick coronal sections containing the somatosensory barrel cortex were pre-incubated in blocking solution 10% bovine serum albumin (BSA), 0.1% sodium azide and 0.02% saponine prepared in TBS (pH 7.4) for 30 min at RT. Then, sections were incubated with a goat anti-CB_1_ receptor antibody (Frontier Institute Co., ltd; goat polyclonal; CB_1_-Go-Af450; FR100610, 1:100) and a guinea pig polyclonal anti-GLAST antibody (Frontier Institute Co., ltd; guinea pig polyclonal; GLAST-GP-Af1000; FR102170, 1:1,000) diluted in 10% BSA/TBS containing 0.1% sodium azide and 0.004% saponine on a shaker for 2 days at 4°C. Sections were incubated with 1.4 nm gold-conjugated rabbit anti-goat pig IgG antibody (Fab fragment, 1:100, #2006, Nanoprobes, Inc., Yaphank, NY, USA) and biotinylated donkey anti-guinea pig IgG antibody (1:200, 706-065-148, Jackson Immuno Research) diluted in 1% BSA/TBS 1x with 0.004% saponin on a shaker for 4 h at RT. Tissues were washed in 1% BSA/TBS on a shaker at RT and incubated with ABC (1:50) prepared in washing solution for 1.5 h at RT. Sections were rinsed with 1% BSA/TBS, stored overnight at 4°C, and post-fixed with 1% glutaraldehyde in TBS for 12 min at RT. After rinsing in double distilled water, gold particles were silver-intensified with the HQ Silver kit (#2012, Nanoprobes, Inc., Yaphank, NY, USA) in the dark for 12 min at RT. The sections were then washed with double distilled water and 0.1 M PB (pH 7.4) for 30 minutes. The biotinylated antibody was revealed with 0.05% DAB in 0.1 M PB (pH 7.4) containing 0.5% Triton X-100 and 0.01% hydrogen peroxide for 3.5 min at RT, followed by washes in 0.1 M PB (pH 7.4). Osmication was done with 1% osmium tetroxide in 0.1 M PB (pH 7.4) in the dark for 20 min. Sections were then washed, dehydrated in graded ethanol, cleared in propylene oxide, pre-embedded in a 1:1 mix of propylene oxide/Epon 812 resin overnight at RT, and finally embedded in pure Epon 812 resin. Electron micrographs were taken with a Hamamatsu FLASH digital camera inserted in a transmission electron microscope (JEOL JEM 1400 Plus). Mitochondria number and morphology and CB_1_ receptor particles were analyzed in 50 pictures at a 4,000 magnification per mouse using Fiji Image J (Schindelin et al., 2012).

### Data collection and statistical analyses

No statistical methods were used to pre-determine sample sizes but they are similar to those reported in previous publications. Experimenters were always blinded to mice genotype but not to treatments. Statistical analyses were performed using GraphPad Prism 10 for Windows (GraphPad Software Inc). Summary results are presented as mean of independent data points ± SEM that represent the number of animals, cultures or astrocytes tested. Datasets were initially tested for normal distribution with the Shapiro-Wilk test and differences between groups were determined by two-tailed Student *t*-test, Mann-Whitney test, Wilcoxon matched-pairs signed-ranks test or two-way ANOVA followed by Šídák’s test for multiple comparisons. Differences were considered to be significant when *p* < 0.05.

## Results

### Autoimmune inflammation disrupts astrocyte mitochondrial dynamics and assembly of respiratory complexes

In order to investigate astrocyte mitochondrial alterations associated to MS we used the chronic EAE model of demyelinating disease in which astrocyte-neuron network dysfunctions emerge as a hallmark of cortical inflammation (Baraibar et al., 2024; Moreno-García et al., 2025). First, we performed immunoelectron microscopy to detect these organelles in astrocyte profiles immunostained for the glutamate aspartate transporter (GLAST). We observed that mitochondrial density within immunoperoxidase-stained cortical astrocyte profiles was significantly increased in EAE mice at acute stages of the disease (**Fig. 1a-b**). Furthermore, quantitative analysis of astrocyte mitochondrial morphology in the EAE cortex revealed significant reductions in the size of these organelles, accompanied by an increase in circularity (**Fig. 1a, c-d**). These anatomical observations suggest that astrocyte activation during autoimmune inflammation involves adaptations in mitochondrial dynamics. At a functional level, preliminary analysis of respiratory efficiency by BNGE followed by IGA showed reduced activity of complex I in cortical mitochondria at acute EAE disease (**Fig. S1a-b**). This result resembles previous observations in demyelinating lesions from MS patients (Dutta et al., 2006). Based on the reported relevance of respiratory SCs for respiration efficiency (Lapuente-Brun et al., 2013; Lopez-Fabuel et al., 2016) we next interrogated the structural organization of the MRC by western blot analysis of complex I subunit NDUFS1 and complex III subunit UQCRC2 in mitochondria purified from cortical tissue (**Fig. S1c-d**). Interestingly, we observed a decrease in the proportions of complex I (**Fig. S1d**, *top panel*) and complex III (**Fig. S1d**, *bottom panel*) that are assembled into SCs relative to those that are free in cortical mitochondria from EAE mice. The protein levels of complex I or III when normalized against the complex II subunit SDHA were similar between control and immunized mice, indicating unaltered expression of mitochondrial complex subunits (**Fig. S1d,** *right panels*). We next sought for changes in the structural organization of the MRC in astrocytes isolated by magnetic cell sorting (**Fig. 1e**). Notably, astrocyte mitochondria from EAE mice showed lower proportions of complex I (**Fig. 1f**, *top panel*) and complex III (**Fig. 1f**, *bottom panel*) assembled to SCs as compared to non-immunized animals, without significant modulation of total complex expression (**Fig. 1f**, *right panels*). Thus, structural remodeling of the MRC correlates to changes in astrocyte mitochondrial numbers and morphology during cortical autoimmune inflammation.

**Figure 1.**
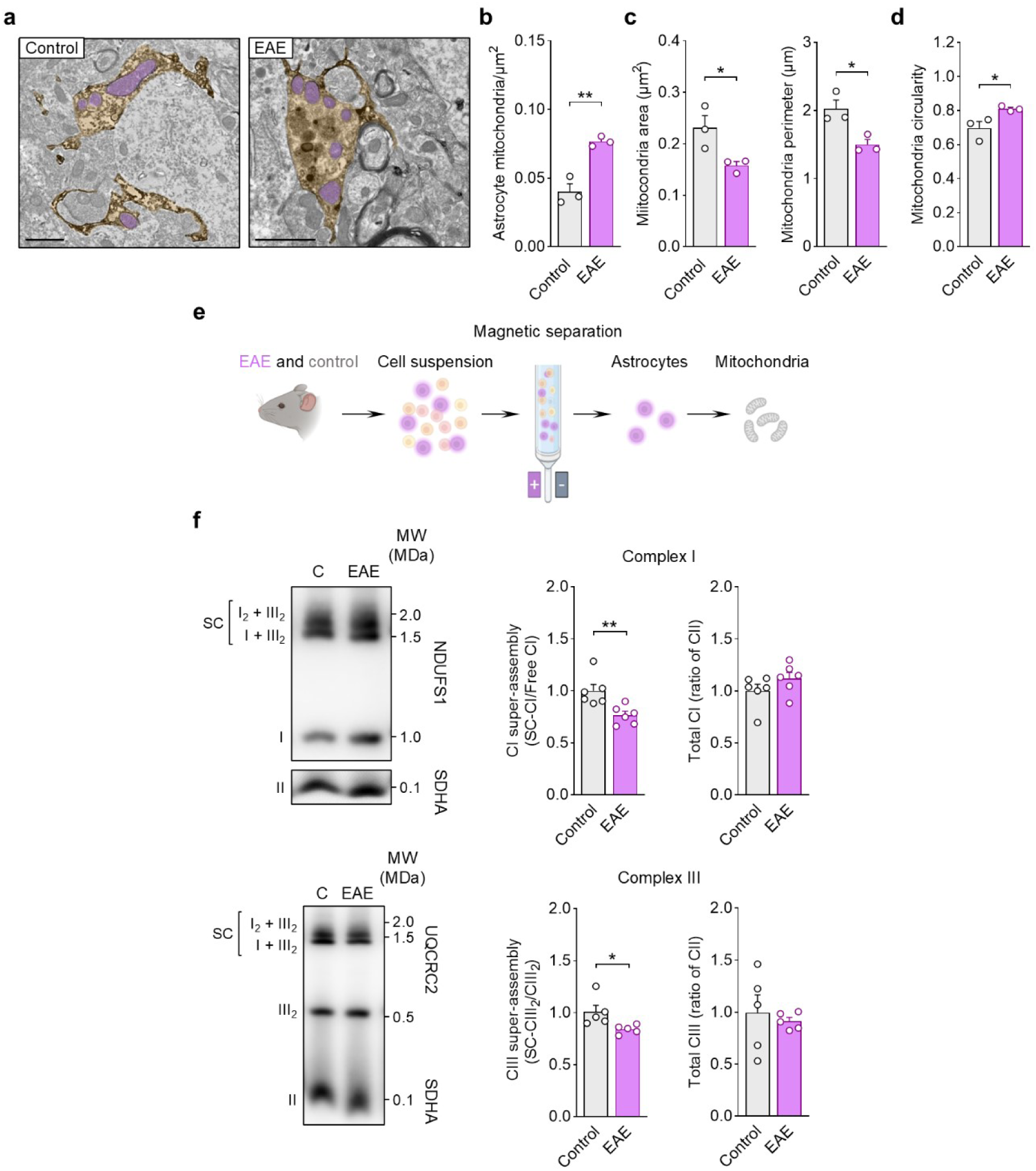
Remodeling of astrocyte mitochondrial networks and respiratory complex assembly during EAE. (**a-d**) Electron microscopy analysis of mitochondria numbers and morphology in the EAE cortex (20 dpi; *n* = 3 mice; score = 4.42 ± 0.30). (**a**) Representative EM images of mitochondria in immunoperoxidase-stained GLAST^+^ astrocyte profiles from control and EAE mice. Scale bars = 1 μm. (**b-d**) Quantitative analysis of astrocyte mitochondria density (**b**), size (**c**) and circularity (**d**) from EM images of control and EAE conditions. (**e**, **f**) *Ex vivo* analysis of mitochondrial respiratory complex assembly in astrocytes purified from EAE mice at acute disease (16 dpi; *n* = 6 mice; score = 3.583 ± 0.15). Digitonin-solubilized mitochondria from astrocytes purified by MACS (**e**) were subjected to BNGE electrophoresis and immunoblotted for the complex I subunit NDUFS1 (**f**, *top panel*) or the complex III subunit UQCRC2 (**f**, *bottom panel*) and the complex II subunit SDHA. The proportion of assembled complex I and complex III was calculated by normalizing SC density values to values corresponding to free NDUFS1 and UQCRC2, respectively. Total expression levels of NDUFS1 and UQCRC2 were normalized to the complex II subunit SDHA (*right panels* in **f**). \**p* < 0.05 and \*\**p* < 0.01, unpaired *t*-test or Mann-Whitney test.

### Inflammatory astrocyte activation impairs mitochondrial function and complex assembly

To strengthen the link between mitochondrial deregulation and astrocyte activation by autoimmune inflammation, we interrogated the metabolic profile of astroglial cells using an *in vitro* system that reproduces, albeit partially, signaling abnormalities of cortical astrocytes in EAE disease (Baraibar et al., 2024). RNA sequencing of purified astrocyte cultures stimulated with the proinflammatory factors TNFα, IL-1α and C1q under serum-free conditions highlighted 370 differentially expressed genes (DEGs) with the majority being upregulated (**Fig. 2a**) (**Table 1**). Confirming the validity of this approach, several genes previously described as induced in astrocytes acutely isolated from EAE mice and serum-free culture models for astrocyte activation (Liddelow et al., 2017; Rothhammer et al., 2016) were also induced in cells stimulated with proinflammatory cytokines.

**Figure 2.**
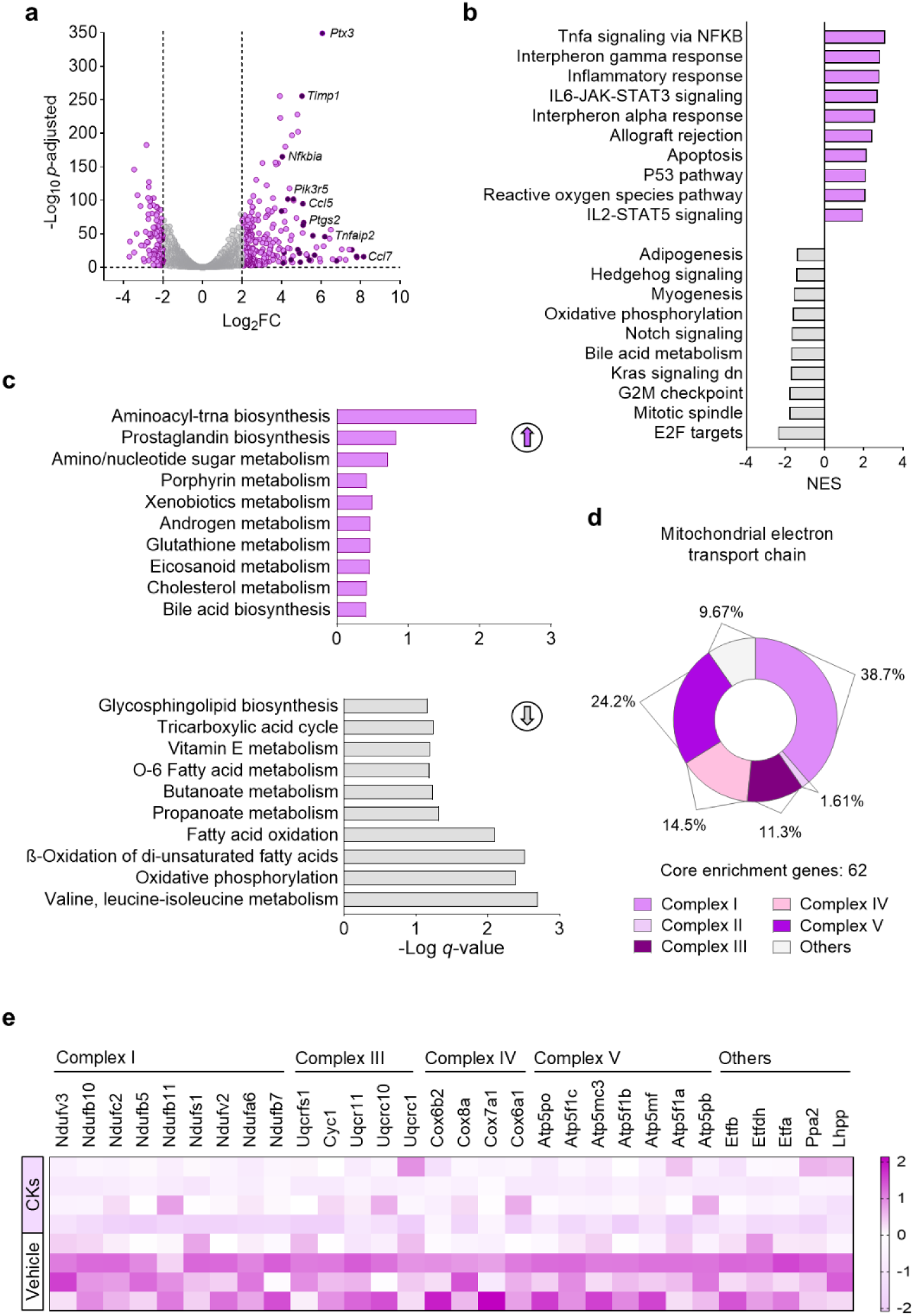
Pathways affected by inflammatory activation of purified astrocyte cultures. Astrocytes were stimulated with TNFα (25 ng/mL), IL-1α (3 ng/mL) and C1q (400 ng/mL) (CKs) for 24 h before transcriptomic analysis (*n* = 4 cultures). (**a**) Volcano plot of DEGs between astrocyte cultures treated with CKs and vehicle-treated cells. X-axis corresponds to log_2_ fold change (FC) and y-axis corresponds to −log_10_ adjusted *p*-value. Purple dots represent the transcripts with an absolute log_2_ FC > 2 and *p*-adjusted < 0.05. Dark purple dots represent the genes belonging to the 3 top upregulated biological processes as depicted in (**b**) among the 60 most upregulated DEGs. Grey dots represent transcripts that do not pass above thresholds. DESeq2 reports an adjusted *p*-value of 0 for *Ptx3* because the true value is below the smallest number representable in double-precision floating point (∼10⁻³⁰⁸). The value shown was recalculated from the DESeq2 Wald statistic (z = 40.25) on a logarithmic scale (raw *p*-value = 3.5 × 10⁻³⁵⁴) and adjusted by the Benjamini–Hochberg procedure over all 21,164 genes with an adjusted *p*-value (padj = 7.4 × 10⁻³⁵⁰). (**b**) Hallmark pathway analysis of RNA sequencing data with GSEA showing 10 top biological processes upregulated and downregulated in astrocyte cultures stimulated with CKs. Gene sets are ordered by normalized enrichment score (NES). For all of them, *p* < 0.01 (Fisher’s exact test). (**c**) GSEA with genome-scale metabolic models (GEMs) showing top metabolic signatures enriched in astrocytes activated with CKs. Positive (pink) and negative (grey) enrichments are indicated with upward- and downward-pointing arrows, respectively. For all of them, *p* < 0.01 (Fisher’s exact test). (**d**) Representative graph showing the proportions of core-enriched genes related to MRC complexes within the OxPhos gene set according to GSEA. (**e**) Heatmap showing the expression profile of the top 30 leading-edge genes in the downregulated OxPhos gene set.

**Table 1.**
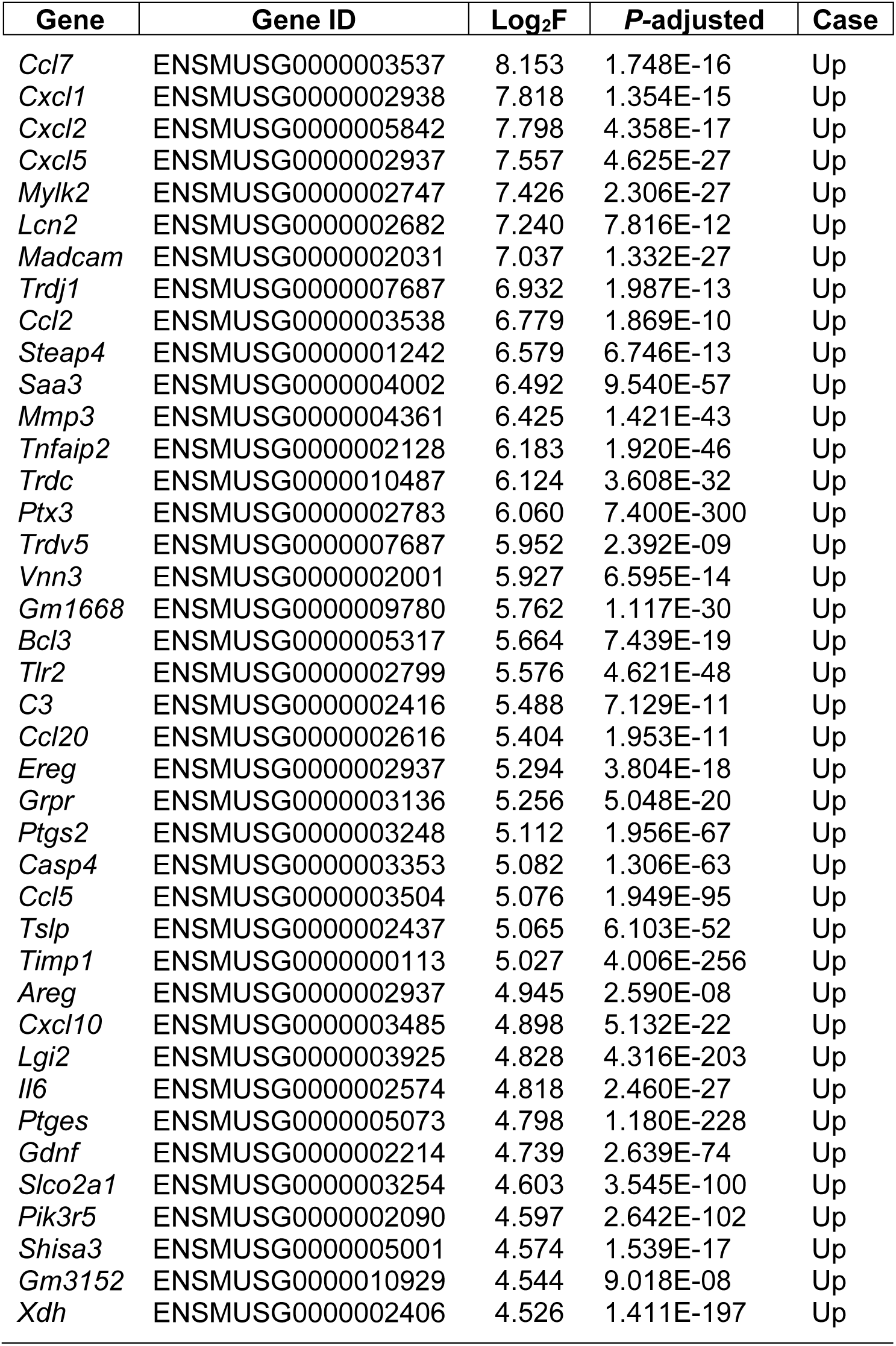
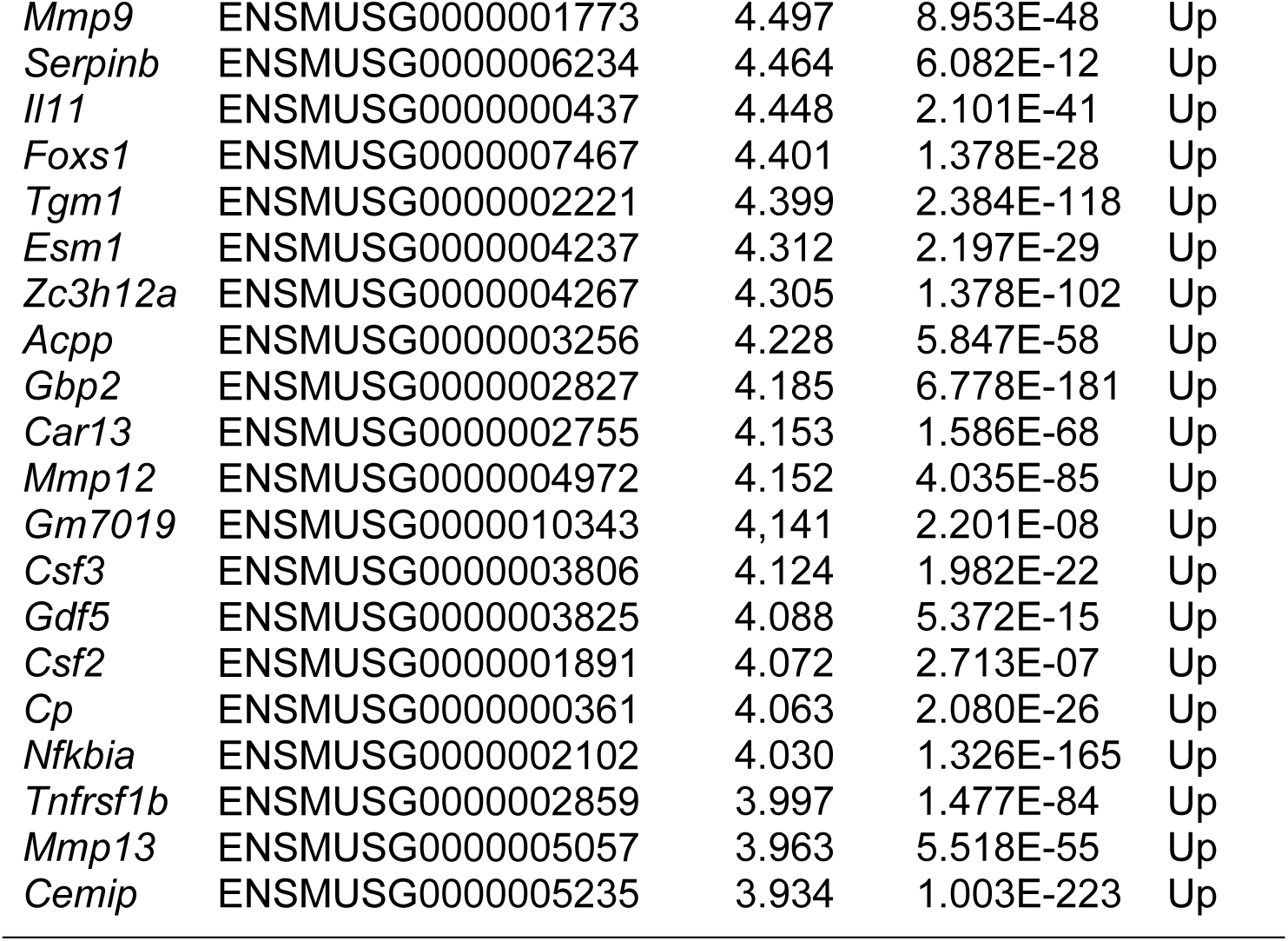
Top 60 DEG genes in astrocyte cultures stimulated with proinflammatory cytokines.

Hallmark pathway analysis of RNA sequencing data with GSEA showed upregulation of gene signatures related to inflammation/immunity and downregulation of genes related to cellular growth, proliferation, differentiation and metabolism (**Fig. 2b**). GSEA of metabolic signatures with GEMs revealed marked gene enrichment in protein and prostaglandin biosynthesis (upregulated) and branched-chain amino acid metabolism, OxPhos and fatty acid oxidation (downregulated) (**Fig. 2c**). The OxPhos gene dataset included 62 core-enriched genes encoding mainly mitochondrial complex I (38.7%), complex V (24.2%), complex IV (14.5%) and complex III (11.3%) showed high overlap in the downregulated transcriptome of astrocytes activated with cytokines (FDR *q*-value < 0.005, GSEA statistical test) (**Fig. 2d-e**). Therefore, induction of an inflammatory phenotype in purified astrocyte cultures downregulates gene signatures related to the MRC among other metabolic patterns.

We next studied the effects of astrocyte activation on mitochondrial function using real-time extracellular flux analysis. Inflammatory stimulation of purified astrocyte cultures did not modulate basal and ATP-coupled respiration (**Fig. 3a-b**). However, we observed a decrease in maximal respiration and spare respiratory capacity indicative of OxPhos impairments in astrocytes activated in a proinflammatory milieu (**Fig. 3a-b**). Digitonin-solubilized mitochondria from stimulated astrocyte cultures showed a trend towards reduced SC assembly following immunoblotting against the complex I subunit NDUFS1 (**Fig. 3c**, *left panel*) and a significant reduction in the proportion of complex III forming SCs (**Fig. 3c**, *right panel*). Changes in SCs formation were not due to deregulated protein levels of complex I or III as determined when normalized against the complex II subunit (**Fig. 3c**). Thus, inflammatory activation of astrocytes *in vitro* translates into changes in the structural organization of the MRC that encompass subtle mitochondrial respiration defects.

**Figure 3.**
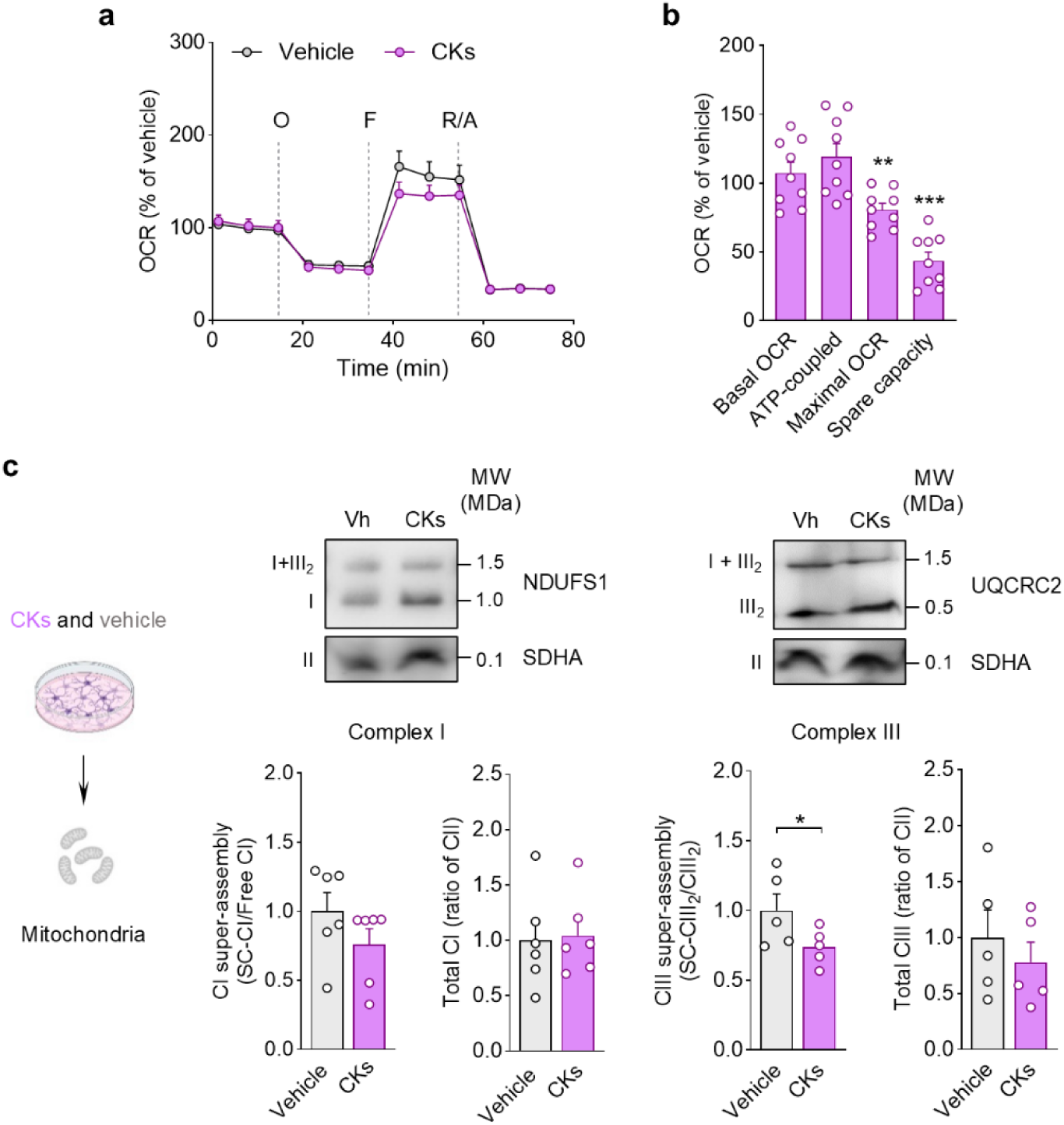
Mitochondria respiration defects and MRC disorganization in astrocytes activated *in vitro*. (**a-b**) Seahorse analysis of mitochondrial respiration in astrocyte cultures using the Mito Stress Test. Basal OCR measurements were performed in astrocytes stimulated with proinflammatory cytokines for 24 h and vehicle-treated cells (*n* = 10 cultures). OCR values in inflammatory astrocytes were expressed as percentages of values in the vehicle condition. Oligomycin (O), FCCP (F), rotenone (R) and antimycin A (A) were added as indicated. (**c**) Analysis of mitochondrial respiratory complex assembly in inflammatory astrocytes activated *in vitro* (*n* = 5-7 cultures). Digitonin-solubilized mitochondria from purified astrocytes cultures were immunoblotted for NDUFS1 (*left panel*) or UQCRC2 (*right panel*) and SDHA. \**p* < 0.05, \*\**p* < 0.01 and \*\*\**p* < 0.001, paired *t*-test or Wilcoxon matched-pairs signed rank test.

### Inflammation alters activity-dependent astrocyte lactate efflux

Analysis of glycolysis-derived proton efflux rates in purified astrocytes revealed increased basal glycolytic flux and reduced glycolytic reserve associated to cell activation with proinflammatory cytokines (**Fig. S2a-b**). Consistent with an improved glycolytic performance, inflammatory astrocytes exhibited higher extracellular lactate concentrations in steady-state conditions (**Fig. S2c**). These combined observations show that inflammation-induced astrocyte activation biases baseline astrocyte metabolism towards glycolysis, but do not provide evidence for deregulated lactate supply in response to neuronal stimuli. Of note, monotypic astrocyte cultures display a limited utility for the analysis of astrocyte-neuron communication mechanisms, as the molecular profile and functional properties of astroglial cells are profoundly influenced by neuronal activity (Hasel et al., 2017; Lange et al., 2012; Mamczur et al., 2015; Ngoc et al., 2025). Thus, we next sought to investigate inflammation-induced adaptations in astrocyte lactate dynamics using mixed cortical cultures of glial and neuronal cells, which represent a validated model of astrocyte metabolic responses to neuronal signals (Lerchundi et al., 2015; Sotelo-Hitschfeld et al., 2015). Cultures expressing the intracellular lactate reporter Laconic were stimulated with proinflammatory cytokines or vehicle and imaged using FRET microscopy (**Fig. 4a**). Consistent with previous reports (Sotelo-Hitschfeld et al., 2015), exposure to high extracellular K^+^ concentrations to simulate neuronal activity led to depletion of the cytosolic astrocyte lactate reservoir (**Fig. 4b-c**). Conversely, astrocytes in mixed cortical cultures responded to near physiological levels of extracellular NH_4_^+^ with an acute rise in cytosolic lactate that results from pyruvate shunting (**Fig. 4b-c**) (Lerchundi et al., 2015). Noteworthily, extracellular K^+^ rises led to smaller responses in inflammatory astrocytes as compared to vehicle-treated cells (**Fig. 4c**, *left panel*) and cytosolic lactate increases induced by extracellular NH_4_^+^ were equally reduced (**Fig. 4c**, *right panel*). Further analysis of Laconic expressing cultures showed that astrocytes stimulated with proinflammatory cytokines display significantly reduced intracellular lactate concentrations in resting conditions as compared to vehicle treated cells (**Fig. 4d**). Thus, exposure to high local levels of proinflammatory mediators triggers reductions in steady-state levels of intracellular lactate and impairs astrocyte responses to extracellular stimuli mirroring neuronal activity.

**Figure 4.**
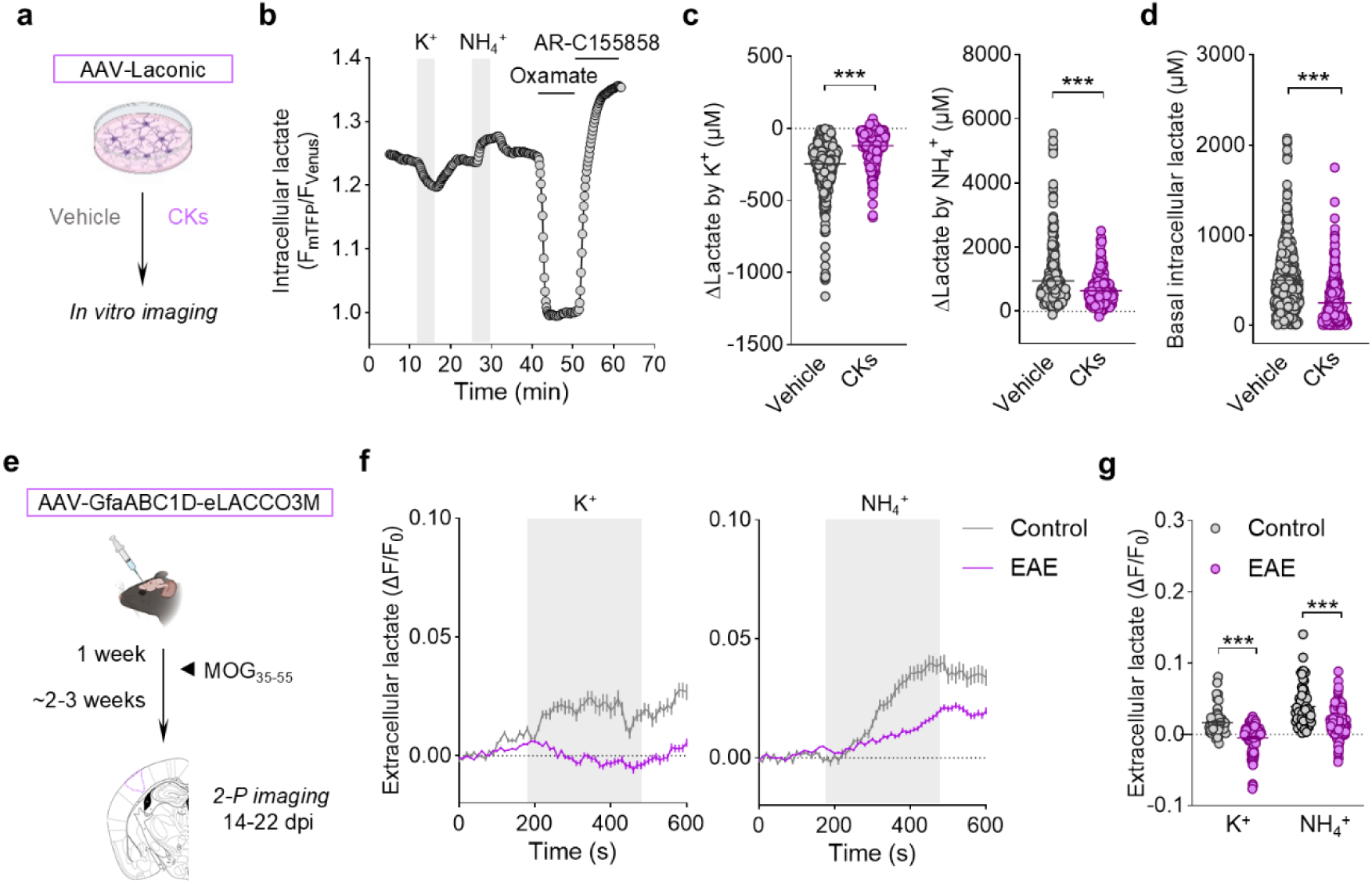
Neuroinflammation impairs activity-dependent astrocyte lactate efflux. (**a**) Imaging of intracellular lactate in astrocyte cultures stimulated with proinflammatory cytokines (CKs). (**b**) Representative time course of intracellular lactate variations by cell superfusion with a high K^+^ (6 mM) solution followed by a NH_4_^+^ (5 mM) after a washout period (*n* = 105 cells). Experiments were calibrated using sodium oxamate (6 mM) and AR-C155858 (1 μM). (**c**) Quantitative analysis shows reduced effect of high K^+^ and NH_4_^+^ to modulate intracellular lactate concentrations in inflammatory astrocytes. (**d**) Resting lactate levels estimated with the protocol in **b** (*n* = 312-489 cells). (**e**) Experimental plan for *ex vivo* recording of extracellular lactate signals in the mouse somatosensory cortex during EAE. Two-photon imaging was performed in acute slices from control and immunized mice previously injected with GfaABC1D-eLACCO3M viral particles. (**f**) Time courses depict the effects of increasing K^+^ (12 mM; *left panel*) (*n* = 5 mice; score = 4.70 ± 0.97) or adding NH_4_^+^ (6 mM, *right panel*) (*n* = 7 mice; score = 5.21 ± 0.75) on the extracellular lactate levels of cortical astrocytes from naive and EAE mice at 14-22 dpi. (**g**) Comparison of normalized responses between experimental groups shows an impaired efficacy of both stimuli to modulate extracellular lactate in the EAE condition (*n* = 56-205 cells). 2-P, two-photon. \*\*\**p* < 0.001; unpaired *t*-test.

To further interrogate the adaptations of astrocyte lactate efflux during CNS inflammation we aimed at measuring extracellular lactate variations in acute slices using the newly developed biosensor eLACCO3M (Liu et al., 2026) infused in the brain cortex. We specifically expressed eLACCO3M in astrocytes using the AAV vector under the control of the GfaABC1D promoter and validated the performance of the fluorescent probe using *ex vivo* two-photon microscopy (**Fig. S3a**). Experiments were performed in the presence of TTX to record lactate dynamics in astrocytes independently of uncontrolled neuronal synaptic activity and thus exclude potential confounding effects from altered excitatory-inhibitory balance associated with cortical EAE pathology (Baraibar et al., 2024; Potter et al., 2016). Imaged in tissue slices, eLACCO3M displayed a rapid boost in the fluorescence signal induced by increasing lactate concentration in the bath solution (**Fig. S3b-c**). To confirm the specificity of these responses, we performed the same experiment using the control vector deLACCO3_ctrol_ (Nasu et al., 2023).

We did not observe significant changes in deLACCO3_ctrol_ upon lactate application (**Fig. S3b-c**). Next, we verified the responses of astrocytic eLACCO3M to stimuli associated to neuronal activity. The extracellular lactate sensor behaved as expected, with an increase in the fluorescence signal on stimulation of astrocyte lactate export by extracellular K^+^ rises and addition of NH_4_^+^ (**Fig. S3d-f**). Lastly, we examined astrocytic lactate dynamics in cortical slices under EAE conditions (**Fig. 4e**). Consistent with our *in vitro* observations (**Fig. 4c**), cortical astrocytes displayed significantly reduced eLACCO3M fluorescence responses to extracellular K^+^ and NH4^+^ rises during the acute phase of EAE as compared to controls (**Fig. 4f-g**). Collectively, these results suggest that astrocyte activation in the context of autoimmune inflammation hinders the ability of these cells to release lactate in response to extracellular stimuli that couple neuronal activity to metabolic supply.

### Impaired coupling of astrocyte CB_1_ receptors to lactate efflux in neuroinflammation

Activation of astrocyte cannabinoid CB_1_ receptors is a crucial mechanism that fine-tunes metabolic coupling and gliotransmission to modulate brain functions in physiological conditions. Canonical CB_1_ receptor signaling in astroglial cells evokes cytosolic calcium elevations and gliotransmitter release while increasing intracellular lactate accumulation and D-serine availability (Eraso-Pichot et al., 2023; Fernández-Moncada et al., 2024; Robin et al., 2018). Conversely, persistent stimulation triggers the activation of CB_1_ receptors located at astrocyte mitochondrial membranes, which results in diminished lactate production eventually leading to neuronal stress in non-inflammatory conditions (Jimenez-Blasco et al., 2020). Of note, previous studies using *in vitro* and *in vivo* methods demonstrate that inflammation hinders calcium responses mediated by astrocyte CB_1_ receptors during cortical EAE pathology (Baraibar et al., 2024). However, whether the adaptations of astrocytes to inflammatory environments involve deregulated metabolic functions mediated by CB_1_ receptors remains uncertain. To explore this possibility, we analyzed the short-term impact of CB_1_ receptor activation in cultured astrocytes expressing the reporter Laconic following 24 h stimulation with proinflammatory cytokines. In good agreement with our previous observations (see Fernández-Moncada et al., 2024), incubation with the cannabinoid agonist WIN55,212-2 induced a rapid increase in Laconic fluorescence that was significantly reduced in inflammatory astrocytes (**Fig. 5a-b**). This *in vitro* finding shows impairments in lactate production stimulated by CB_1_ receptors. We next asked about adaptations in lactate efflux mediated by CB_1_ receptors in an inflammatory tissue context. To explore this possibility, we first investigated whether activation of astrocyte CB_1_ receptors triggers lactate production and release in brain slices using the eLACCO3M biosensor as described in **Fig. S3a**. When the slices were exposed to WIN55,212-2, cortical astrocytes expressing eLACCO3M responded with an increase in fluorescence intensity indicative of a rise in extracellular lactate (**Fig. 5c-f**). Consistent with CB_1_ receptor-mediated regulation, we found that the eLACCO3M responses to the cannabinoid agonist were prevented by bath perfusion of the selective antagonist AM251 (**Fig. 5d, f**). Furthermore, deletion of the *Cnr1* gene by viral expression of Cre-recombinase in cortical astrocytes of CB_1_^f/f^ mice (aCB_1_-KO) abolished the increase in eLACCO signal induced by WIN55,212-2 (**Fig. 5e, f**). Thus, astrocyte CB_1_ receptors mediate extracellular lactate rises in tissue slices. We next investigated adaptations of cannabinoid-induced lactate efflux in cortical tissue from EAE mice. Mirroring observations in cultures simulated with proinflammatory cytokines (**Fig. 4f-g**), astrocytes within the brain cortex of mice at acute EAE disease displayed significantly reduced eLACCO responses to WIN55,212-2 (**Fig. 5g-h**). Altogether, our results in cultured cells and tissue slices suggest that inflammation impairs the ability of astrocyte CB_1_ receptor agonism to promote lactate production and release.

**Figure 5.**
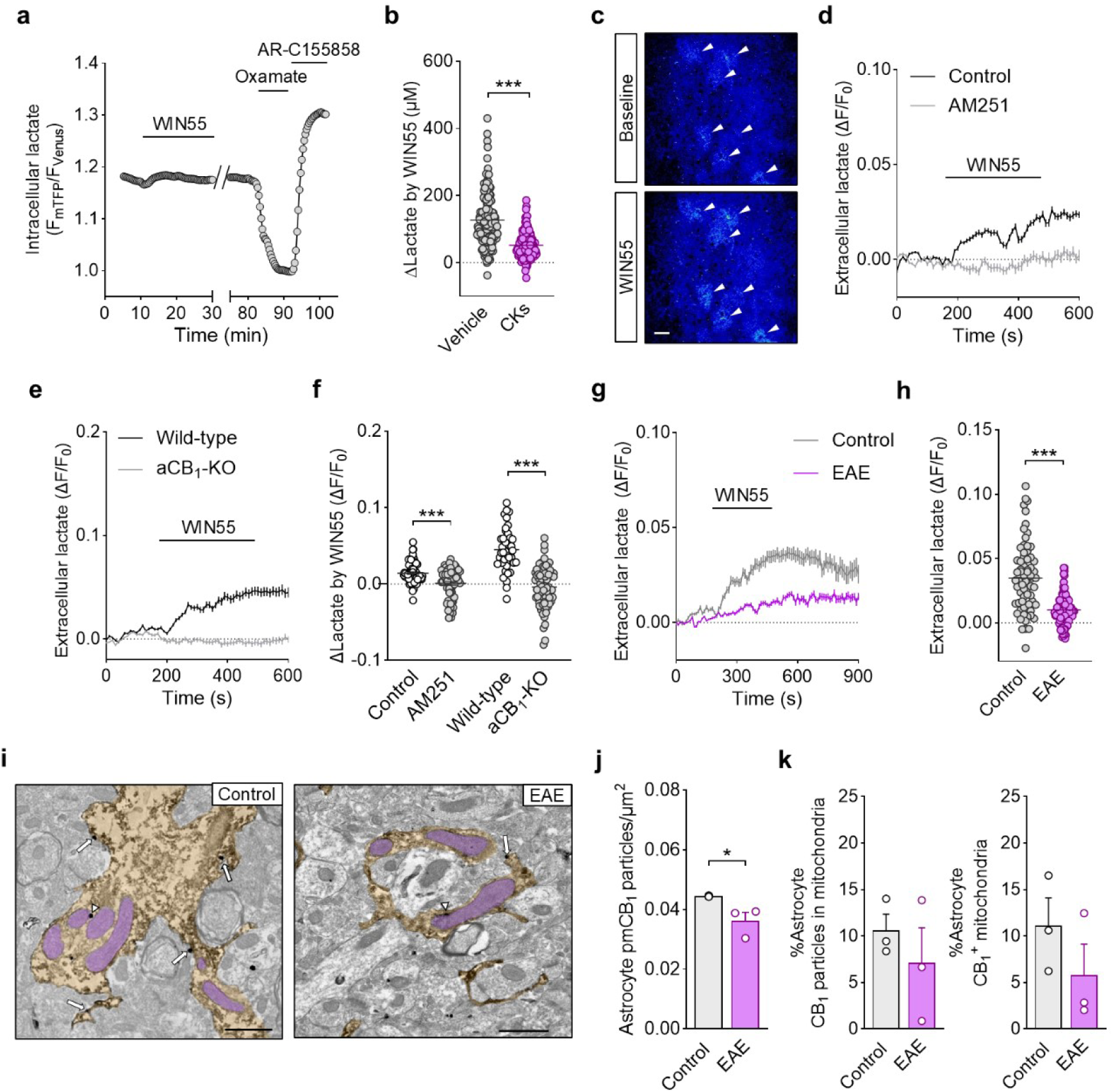
Inflammation-induced defects in CB_1_ receptor-mediated astrocyte lactate production and release. (**a-b**) Analysis of astrocyte lactate responses to the activation of CB_1_ receptors *in vitro*. Representative time-course (**a**) and quantitative analysis (**b**) of intracellular lactate variations by the cannabinoid agonist WIN55,212-2 (WIN55; 1 μM) in astrocyte cultures stimulated with proinflammatory cytokines (CKs) (*n* = 141-158 cells). (**c-f**) CB_1_ receptors mediate astrocyte lactate responses in acute cortical slices. (**c**) Pseudocolor images of eLACCO3M fluorescence in cortical tissue before and after bath application of WIN55 (3 µM). Scale bar = 50 µm. (**d-f**) Comparative analysis of extracellular lactate variations induced by WIN55 alone and in the presence of the CB_1_ receptor antagonist AM251 (3 µM; *n* = 70-84 cells) (**d**), or in slices from CB_1_^f/f^ mice injected with AAV9-GFAP-Cre (aCB_1_-KO; *n* = 43-77 cells) (**e**). (**g-h**) Effect of WIN55 on extracellular lactate measurements in EAE mice and naïve controls (14-22 dpi; *n* = 5 mice; score = 5.40 ± 0.93). Data points correspond to *n* = 70-107 cells. (**i-k**) Localization of CB_1_ receptors and GLAST in the somatosensory cortex layer V-VI of naïve and EAE mice (*n* = 3 mice). Representative micrographs depict immunopositive astrocyte processes (orange) containing mitochondria (purple) in both experimental groups. CB_1_ particles were detected in close association to the astrocyte plasmatic (pmCB_1_; arrows) and mitochondrial (arrowheads) membranes. Scale bars = 1 µm. (**j**, **k**) Statistical assessment of immunoparticle distribution showed reductions of CB_1_ receptor density in the astrocyte plasma membrane (**j**) and unaltered expression levels in the mitochondrial compartment (**k**). \*\*\**p* < 0.001 and \**p* < 0.05; unpaired *t*-test or Mann-Whitney test.

Next, we interrogated the mechanisms underlying the observed deficits in lactate responses to cannabinoid agonists. According to previous studies in the EAE model, impairments in CB_1_ receptor-mediated astrocyte responses associated to neuroinflammation may result from defects in intracellular pathways operated by G proteins as well as from reduced expression levels of the receptor protein (Baraibar et al., 2024). Indeed, astrocytes activated during EAE pathology display reduced transcript levels of CB_1_ receptors according to gene expression studies in cells purified from the mouse forebrain (Moreno-García et al., 2020). To disentangle this specific point, we investigated the astrocytic expression of CB_1_ receptors in the somatosensory cortex of EAE mice using high-resolution immunoelectron microscopy. Astrocytes and their processes were identified by DAB immunodeposits of GLAST and the CB_1_ receptor was detected in parallel by immunogold labelling. As expected from previous localization studies throughout the mouse forebrain (Gutiérrez-Rodríguez et al., 2018; Gómez-Sotres et al., 2024; Jimenez-Blasco et al., 2020), we detected DAB immunopositive astrocytic processes in the barrel somatosensory cortex that displayed CB_1_ receptor immunoparticles in close juxtaposition to both the plasma membrane and the mitochondrial membranes (**Fig. 5i**). Notably, the density of CB_1_ receptor immunoparticles on astrocytic plasma membranes was significantly reduced in EAE mice during acute disease (**Fig. 5j**). By contrast, the proportion of astrocyte CB_1_ receptor immunoparticles located on mitochondrial membranes within the EAE cortex did not differ significantly to that measured in the control group (**Fig. 5k**, *left panel*). Similarly, the percentage of CB_1_ receptor immunopositive mitochondria within cortical astrocyte profiles was similar between naïve and EAE conditions (**Fig. 5k**, *right panel*). Thus, cortical astrocytes exhibit a selective reduction in the expression levels of CB_1_ receptors at the plasma membrane that may underlie, at least in part, impairments in lactate efflux during EAE.

## Discussion

The present study offers a combined analysis of astrocyte bioenergetic adaptations in autoimmune inflammation targeting mitochondria and lactate surge. Key findings include: (1) disturbances in the structural organization of the MRC encompass changes in astrocyte mitochondrial dynamics during EAE; (2) remodeling of astrocyte mitochondrial networks during cortical inflammation occurs in concert with deficits in lactate release responses to extracellular stimuli that mirror neuronal activity, and (3) astrocyte activation is associated with reductions in the plasma levels of CB_1_R in the inflamed cortex. These results suggest complex metabolic adaptations involving astrocyte mitochondria and lactate production/release machinery with implications on our understanding of cortical pathology in MS.

The investigation of mitochondrial dysfunction in MS using *postmortem* tissue and preclinical models shows altered dynamics, structure and function of these organelles due to disease-induced changes in energy demand and production (Dutta et al., 2006; Licht-Mayer et al., 2020; Witte et al., 2009). These converging lines of evidence, predominantly focused in neurons, support the hypothesis that defective mitochondria contribute to neurodegeneration and irreversible functional impairment in MS (Atkinson et al., 2023). Strikingly, a paucity of studies has examined the astrocyte-specific mitochondrial pathophysiology of MS despite the fundamental bioenergetic roles of these cells in nervous tissue and their critical contribution to the disease progression (Bolaños & Magistretti, 2025; Ponath et al., 2018). As far as we are aware, the present study is the first report of changes in mitochondrial networks affecting cortical astrocytes in preclinical MS. Our quantitative electron microscopy analysis highlights an increased mitochondrial density in astrocyte profiles within the EAE cortex that sits well with previous histopathological observations in active MS lesions (Witte et al., 2009). Ultrastructural examination further unveiled reductions in astrocyte mitochondrial size associated with an increased circularity. These unprecedented observations suggest a disruption of astrocyte mitochondrial dynamics towards fragmentation with potential consequences in the pathophysiology of cortical MS (Atkinson et al., 2023; Göbel et al., 2018; Youle & van der Bliek, 2012). The initiating insult that triggers the reorganization of astrocyte mitochondrial networks during EAE remains unclear, but current evidence points to the involvement of the local inflammatory microenvironment. Inflammation driven by peripheral and innate immune cells is a feature of cortical pathology induced by immunization with MOG (Baraibar et al., 2024; Colomer et al., 2026; Potter et al., 2016) and astrocytes within the brain cortex react to acute inflammatory injury by increasing mitochondrial fission (Motori et al., 2013). Further characterization in culture systems and acute slices has demonstrated that cortical astrocytes increase the activation of the pro-fission protein DRP1 as key signaling event that leads to mitochondria fragmentation in response to inflammatory stimuli (Motori et al., 2013). While the present study did not address the specific molecular mechanisms leading to changes in mitochondrial dynamics during cortical demyelinating pathology, it is noteworthy that pluripotent stem cell-derived astrocytes from MS patients display increased DRP1 levels and ultrastructural alterations indicative of mitochondrial fragmentations (Ghirotto et al., 2022). Interestingly, the study of Ghirotto et al. (2022) reported an exacerbated production of proinflammatory chemokines associated to upregulated mitochondrial fission in MS astrocytes, thus raising the possibility that aberrant astrocyte-based mitochondrial dynamics participates as pathogenic mechanism of disease progression. Beyond these observations, the consequences of mitochondrial fission on astrocyte-derived inflammation remain scantly known. Independent laboratories have demonstrated that the inhibition of DRP1 function is protective in experimental models of demyelination and neurodegenerative disease, but the observed benefits cannot be solely attributed to changes in mitochondrial division, as additional anti-inflammatory mechanisms may be involved (Cai et al., 2024; Luo et al., 2017). In the aforementioned context, the present anatomical findings, that establish a solid link between cortical autoimmune demyelinating pathology and astrocyte mitochondrial network deregulation, warrant further analysis of the mechanisms and implications of astroglial mitochondrial reshaping in MS.

Over the last decade, a number of studies have established associations between MRC disorganization and mitochondrial dysfunction in neurodegenerative disease. The strongest evidence comes from neurological disorders in which disease-related genes are linked to mitochondrial pathways such as Parkinsońs disease or Friedreich ataxia (Doni et al., 2021; Hara et al., 2024; Heo et al., 2012; Lopez-Fabuel et al., 2017). However, experimental support for this same association in MS is currently lacking. Here we report that, in cortical tissue and astrocyte-enriched preparations, complex I and complex III are disassembled from respiratory SCs during EAE pathology. Thus, remodeling of the MRC encompasses aberrant astrocyte mitochondrial dynamics during autoimmune inflammation, further supporting the intimate link between ultrastructural features that critically shape oxidative metabolism and respiratory efficiency (Baker et al., 2019). Decreased CI interaction with CIII determines low electron transfer through the MRC and high generation of mROS in astrocytes (Lopez-Fabuel et al., 2016). However, one limitation of the present study is that it did not address astrocyte-specific mitochondrial function in the EAE model. Conversely, using a culture system we show that cortical astrocytes respond to proinflammatory stimuli with modest reductions in maximal and spare respiratory capacity that are paralleled by a decrease in SC-assembled complex III. Although these results offer limited support for an association between astrocyte SC disassembly and respiration defects in cortical neuroinflammation, the observed adaptations are consistent with the reductions in OxPhos genes measured by transcriptomic analysis in our murine model and mirror previous reports of mitochondrial bioenergetic impairments in rodent and human astrocytes exposed to inflammatory stimuli (Motori et al., 2013; Pamies et al., 2021; Russ et al., 2021). Remarkably, the selective reductions in maximal respiration rates and spare capacity that we report here fully recapitulate the functional deficits that occur in concert with the fragmentation of mitochondrial networks in cortical astrocytes activated *in vitro* (Motori et al., 2013). It should be noted, however, that the bioenergetic reprogramming of cultured astrocytes during inflammatory activation may translate into an increased OxPhos performance according to some studies (Chao et al., 2019; Meyer et al., 2022; Robb et al., 2020). The reasons for these contradictory findings remain uncertain, but they may stem from discrepancies between experimental protocols for the purification, maintenance and activation of astrocyte cultures. In this regard, it is worth mentioning that the metabolic adaptations of astrocytes to stimulation with interferon gamma (IFNγ) and lipopolysaccharide (LPS) are stimulus-dependent and evolve as a function of the exposure time, with short-term and long-term treatments evoking distinct adaptations that affect both OxPhos and glycolysis in culture systems (Motori et al., 2013; Robb et al., 2020). These results underscore the complexity of astrocyte bioenergetic reprogramming during the acquisition of reactive states in inflammatory microenvironments and highlight the need of a comprehensive evaluation using disease-specific model systems and astrocyte-selective assessments in a tissue context.

The activity-dependent supply of lactate as bioenergetic messenger is a distinctive feature of astrocyte physiology that shapes brain function and behavior (Magistretti & Allaman, 2018). Current hypothesis, based on a combination of transcriptomic, immunohistochemical and functional readouts from *in vitro* and *in vivo* models, postulate that defects in astrocyte-neuron lactate shuttle are relevant to neuronal damage in the context of neurodegenerative disease (Bittner et al., 2023; Bolaños & Magistretti, 2025; Cunnane et al., 2020). A counterintuitive but consistent observation, however, is that astrocytes reacting to inflammatory stimuli increase their glycolytic rates (Almeida et al., 2004; McComish et al., 2024; Véga et al., 2002; Xiong et al., 2025). This metabolic switch is interpreted as adaptative strategy that ensures astrocyte performance and survival during the acquisition of reactive states while conferring neuronal protection in neuroinflammatory contexts. However, the majority of these studies do not provide insights on astrocyte lactate responses operated by neuronal stimuli and the metabolic adaptations of astroglial cells to neuroinflammation *in vivo* have not been tested systematically yet. Consistent with the above-mentioned line of evidence, in this study we report a marked increase in glycolytic rates, as judged by extracellular flux analysis and lactate production, in monotypic astrocyte cultures stimulated with proinflammatory cytokines. By contrast, real-time measurements of intracellular lactate in cortical cultures of neurons and glial cells evidenced that inflammatory activation engages reductions in the steady-state astrocyte lactate reservoir. This novel piece of evidence points to a limited capability of astrocytes to mobilize and supply lactate upon neuronal workload. In support of this hypothesis, reactive astrocytes in mixed cortical cultures displayed impaired lactate responses to well-established metabolic recruiters such as K^+^ and NH4^+^ (Barros et al., 2023). These findings align with early evidence of reductions in glutamate-induced metabolic burst in inflammatory astrocytes activated *in vitro* (Gavillet et al., 2008). Our analysis of extracellular lactate dynamics in brain slices from EAE mice corroborated, at the tissue level, impairments in astrocytes metabolic responses to extracellular stimuli, further suggesting that autoimmune inflammation perturbs the ability of cortical astrocyte networks to support neuronal bioenergetic demand. Altogether, these observations cement the hypothesis that maladaptive astrocyte-neuron metabolic coupling in terms of lactate surge is a feature of cortical demyelinating pathology. A remaining question is how local inflammation translates into a general defect in astrocyte bioenergetic feedback to metabolic recruiters that signal over different spatiotemporal domains and implicate different intracellular mechanisms (Barros et al., 2023). Previous studies in the EAE model suggest that inflammatory stimuli dampen the astrocytic production of lactate through cytosolic phospholipase A2-dependent mechanisms that limit hexokinase activity and boost the mitochondrial oxidation of pyruvate (Chao et al., 2019). These immunometabolic adaptations may underlie defects in the ability of reactive astrocytes to supplement the extracellular lactate pool in response to neuronal stimuli such as K^+^ and NH4^+^ (Lerchundi et al., 2015; Sotelo-Hitschfeld et al., 2015), although the implication of alternative molecular mechanisms cannot be discarded. We recently found that autoimmune inflammation dramatically disrupts the calcium signaling machinery of cortical astrocyte networks (Baraibar et al., 2024; Moreno-García et al., 2025), with potential implications in astrocyte-neuron metabolic flux (Cabral-Costa et al., 2022; Horvat et al., 2021). This possibility deserves further investigation. Advances in cellular imaging combining calcium and metabolic indicators, genetic manipulation of metabolic pathways, and optogenetic modulation of neuronal activity, should shed light on the mechanisms and implications of deregulated astrocyte lactate surge in neuroinflammatory disorders such as MS.

Endocannabinoids have recently emerged as novel metabolic messengers that promote astrocyte lactate efflux to modulate synaptic function and cognition through complex mechanisms (Fernández-Moncada et al., 2024). In this study, we confirm previous observations that CB_1_ receptors engage lactate production and release in culture models and corroborate this finding at the tissue-level using a slice-based imaging approach. We also provide unprecedented evidence that autoimmune demyelinating pathology hinders the ability of endocannabinoids to engage astrocyte metabolic processes that lead to lactate supply. Furthermore, our electron microscopy analysis in the EAE model demonstrates that cortical inflammation downregulates the expression of CB_1_ receptors on astrocyte plasmatic membranes. These anatomical results thus provide a mechanistic basis for the observed deficits in lactate efflux (present study), calcium signaling and gliotransmission (Baraibar et al., 2024), that encompass astrocyte activation in the EAE cortex. Interestingly, comparative analysis of CB_1_ receptor expression in different astrocyte compartments highlights that cortical inflammation triggers a selective reduction in the population associated to plasmatic membranes, while the mitochondrial receptor pool, that hampers glycolysis and lactate production (Jimenez-Blasco et al., 2020), remained unaffected. Although these results should be interpreted with caution, given the limited number of mice included in our study, it seems reasonably safe to postulate that the extent and time-course of changes in CB_1_ receptor expression in specific subcellular locations determines, at least in part, the output of astrocyte-mediated endocannabinoid signaling during autoimmune inflammation, in terms of gliotransmission and metabolic functions. These findings, that have implications in MS symptomatology and progression, require further investigation using genetic tools that allow the cell-specific modulation of selective receptor pools in the disease context.

In summary, the present work identifies astrocyte mitochondrial network fragmentation and respiratory complex disassembly as features of autoimmune inflammation and unveil astrocyte bioenergetic failure in supporting neuronal activity within the inflamed cortex. Our results provide plausible evidence of deficits in the astrocyte-neuron metabolic unit as potential pathogenic mechanism of cortical dysfunction in MS and suggest that interventions focused on restoring mitochondrial phenotypes and lactate supply may hold promise to mitigate MS symptomatology and avert disease progression.

## Data and code availability

This study reports new RNA sequencing datasets generated from pure astrocyte cultures that have been deposited at GEO. Raw data from biochemical, electron microscopy and imaging experiments are available upon reasonable request.

## Acknowledgements

We thank all members of the S Matós, G Marsicano’s and BrainStrim laboratories for useful discussions and advice, the personnel from Achucarro Basque Center of Neuroscience Imaging and Cell Analytics Facilities, and the personnel of the Animal Facilities of the University of the Basque Country and Neurocentre Magendie for mouse care. **Figures 1e**, **3c**, **4a**, **4e** and **S1c** were created using BioRender.com.

## Author contributions

I.F.-M., J.P.B., G.M. and S.M. conceived the study. T.C. performed and analyzed MRC assembly experiments. A.M.B. and I.F.-M. conceptualized and performed imaging experiments. N.P., I.B. and P.G. performed and supervised electron microscopy analysis. C.M., A.B.-C. and E.S.-M. performed EAE experiments. N.C.-M. carried out stereotactic injections for *ex vivo* image analysis. G.P. performed RNAseq analysis. D.J.-B. and J.P.B. supervised MRC assembly experiments and analysis. Y.N., A.C.P.Z. and E.S.-G. provided critical reagents and resources. A.M.B., T.C., N.P. and S.M. prepared the figures. S.M. drafted the manuscript. All authors reviewed and commented on the manuscript.

---

**Supplementary Figure 1.**
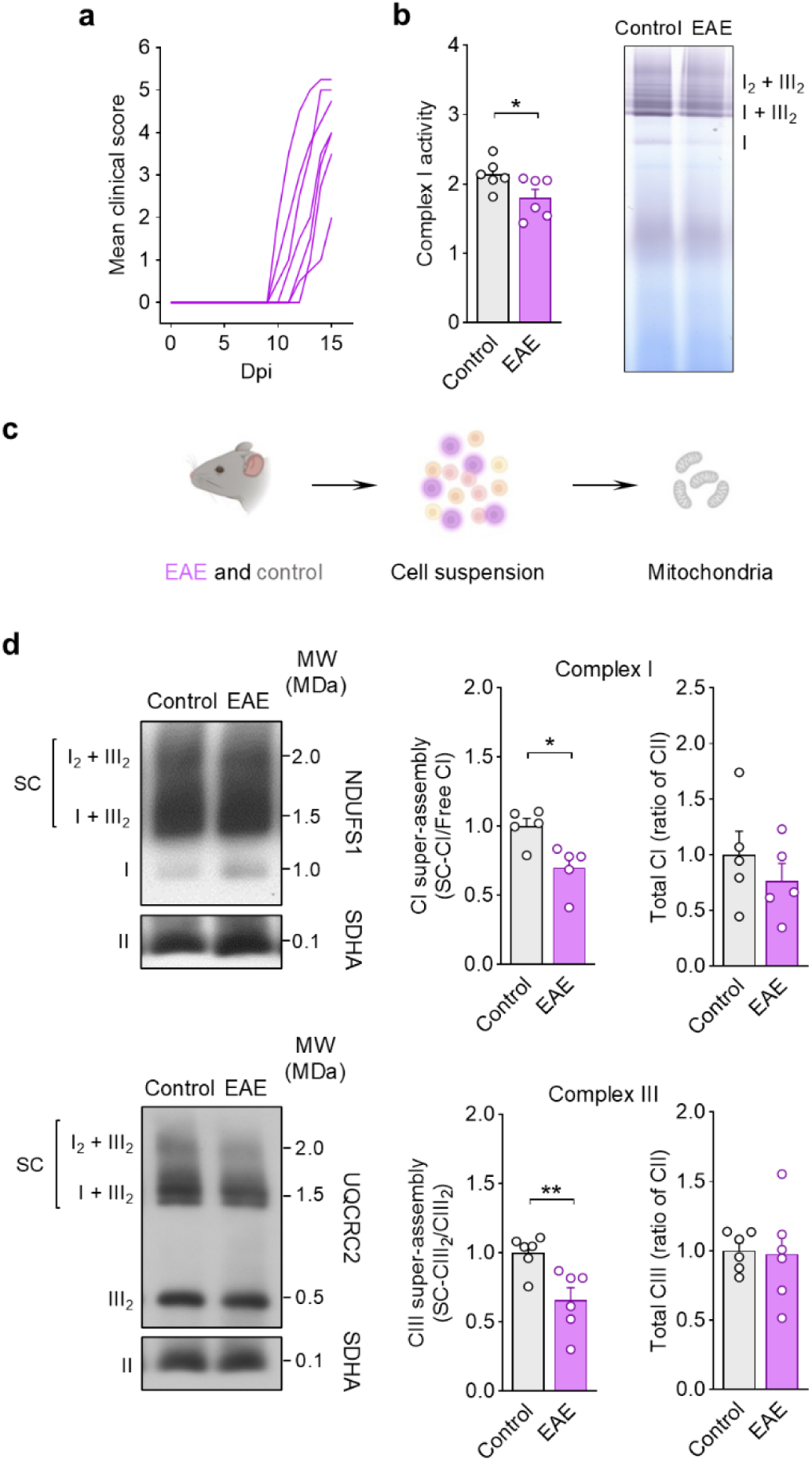
EAE disrupts mitochondrial SC assembly in the brain cortex. (**a**) Clinical scores of EAE mice included in the analysis of MRC supercomplex assembly (15 dpi; *n* = 7 mice). (**b**) IGA of complex I in cortical tissue from control and EAE mice (*n* = 6 mice). (**c-d**) Digitonin-solubilized cortical mitochondria from mice under acute EAE were subjected to BNGE electrophoresis and immunoblotted for the complex I subunit NDUFS1 (**d**, *top panel*) and the complex III subunit UQCRC2 (**d**, *bottom panel*) (*n* = 5-7 mice). Total expression levels of NDUFS1 and UQCRC2 were normalized to the complex II subunit SDHA. Dpi, days post-immunization. \**p* < 0.05, unpaired *t*-test or Mann-Whitney test.

**Supplementary Figure 2.**
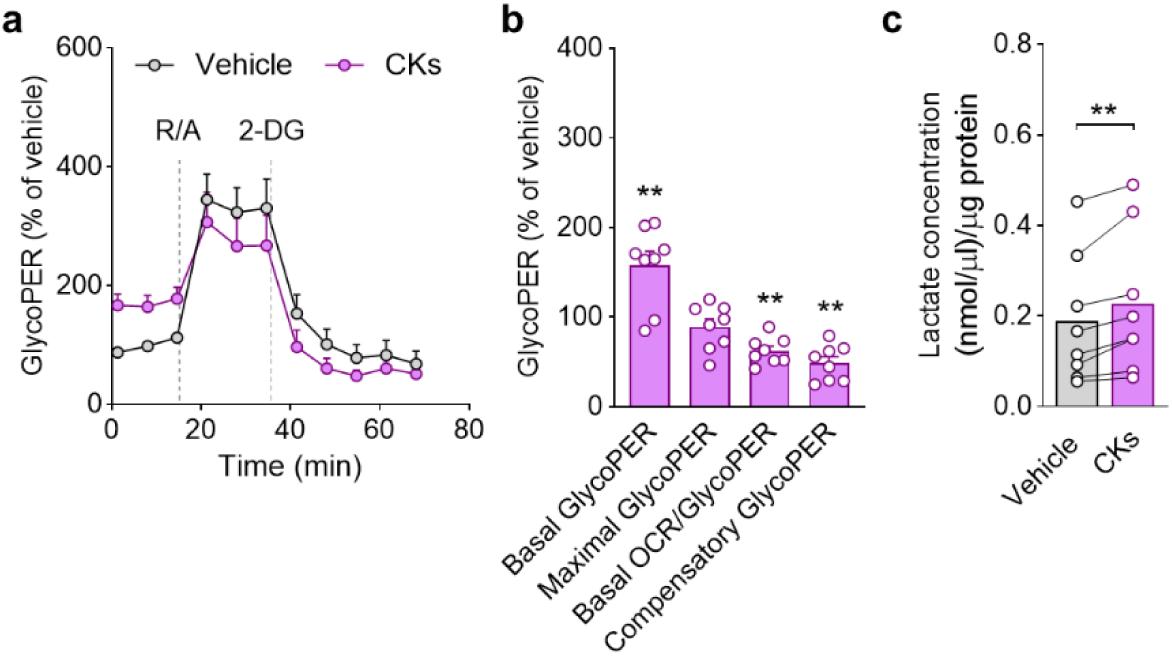
Dysregulated glycolytic flux in inflammatory astrocyte cultures. (**a-b**) Glycolytic flux assessed by GlycoPER in astrocyte cultures stimulated with CKs and vehicle-treated cells (*n* = 8 cultures). GlycoPER values in stimulated cells were expressed as percentages of vehicle condition. Rotenone (R), antimycin A (A) and 2-deoxyglucose (2-DG) were added as indicated. (**c**) Extracellular lactate measurement with an enzymatic kit in control and inflammatory astrocyte cultures (*n* = 8 cultures). \*\**p* < 0.01, paired *t*-test or Wilcoxon matched-pairs signed rank test.

**Supplementary Figure 3.**
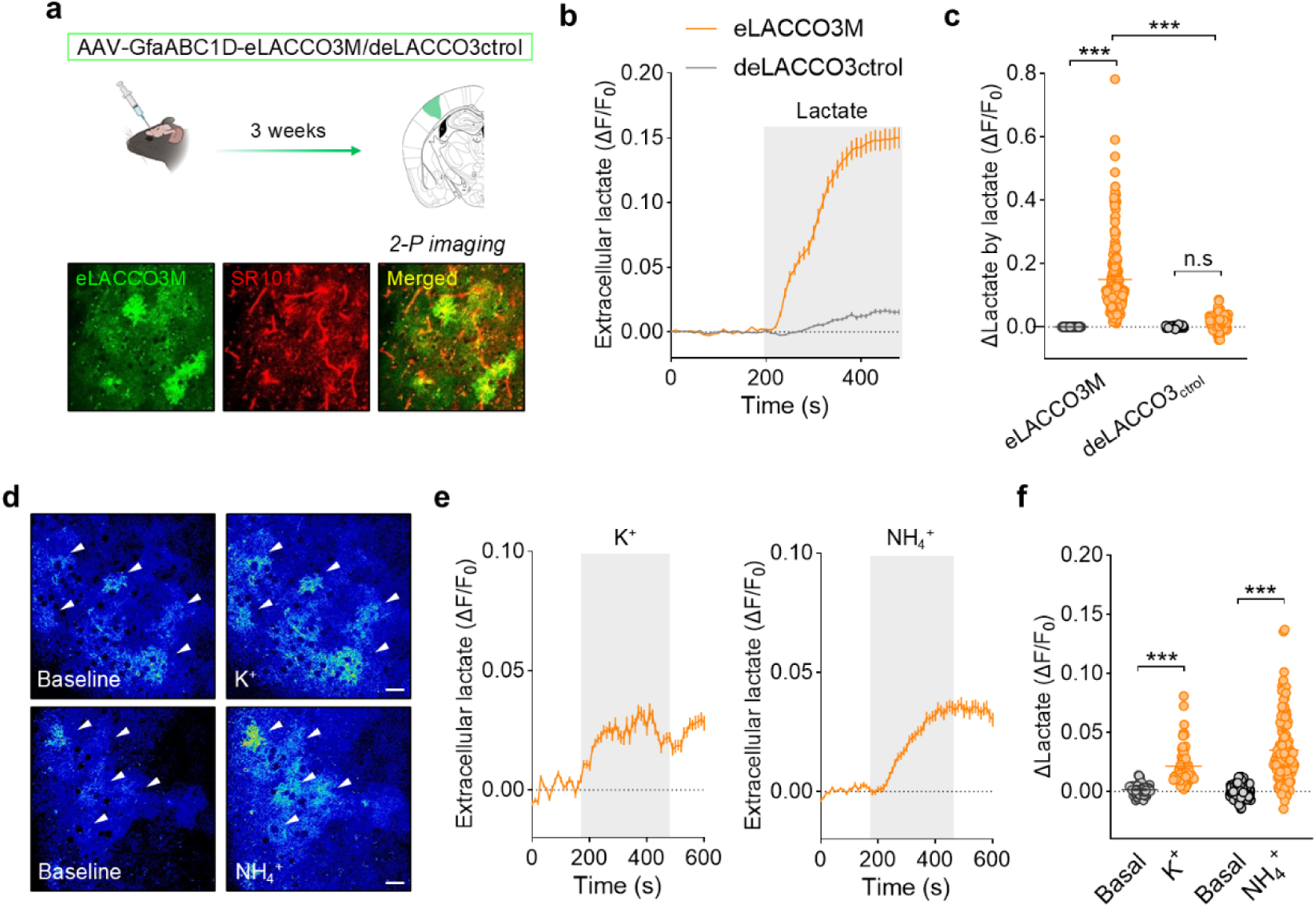
*Ex vivo* determination of extracellular lactate variations in cortical astrocytes. (**a**) Acute slices from mice previously injected with GfaABC1D-eLACCO3M/deLACCO3ctrol viral particles were imaged in a two-photon microscope for measurement of extracellular lactate. Images of eLACCO3M expression in the somatosensory cortex depicting biosensor colocalization with the astrocytic marker sulforhodamine 101 (20 mg/mL; 200 µL i.p. 60 min before sacrifice). Scale bar = 50 µm. (**b-c**) Validation of the eLACCO3M biosensor in acute cortical slices. The fluorescence intensity of eLACCO3M increased upon addition of 10 mM lactate whereas no significant changes were observed in slices from deLACCO3ctrol-injected mice (*n* = 127-205 cells). (**d**) Representative images of cortical eLACCO3M fluorescence signals accumulated over a 1 min period in baseline conditions and following application of high extracellular K^+^ (12 mM; *top panel*) or NH_4_^+^ (6 mM; *bottom panel*). Scale bars = 50 µm. (**e**, **f**) Time courses and dot plots depict extracellular lactate variations measured in acute slices following perfusion of a high K^+^ solution (*n* = 62 cells) or NH4^+^ (*n* = 117 cells). \*\*\**p* < 0.001, two-way ANOVA followed by Šídák’s test for multiple comparisons (**c**) and Wilcoxon matched-pairs signed rank test (**f**).

## Bibliography

Almeida, A., Jimenez-Blasco, D., & Bolaños, J. P. (2023). Cross-talk between energy and redox metabolism in astrocyte-neuron functional cooperation. Essays Biochem, 67(1), 17–26. 10.1042/EBC20220075

Almeida, A., Moncada, S., & Bolaños, J. P. (2004). Nitric oxide switches on glycolysis through the AMP protein kinase and 6-phosphofructo-2-kinase pathway. Nat Cell Biol, 6(1), 45–51. 10.1038/ncb1080

Aten, S., Kiyoshi, C. M., Arzola, E. P., Patterson, J. A., Taylor, A. T., Du, Y.,…Zhou, M. (2022). Ultrastructural view of astrocyte arborization, astrocyte-astrocyte and astrocyte-synapse contacts, intracellular vesicle-like structures, and mitochondrial network. Prog Neurobiol, 213, 102264. 10.1016/j.pneurobio.2022.102264

Atkinson, K. C., Osunde, M., & Tiwari-Woodruff, S. K. (2023). The complexities of investigating mitochondria dynamics in multiple sclerosis and mouse models of MS. Front Neurosci, 17, 1144896. 10.3389/fnins.2023.1144896

Baker, N., Patel, J., & Khacho, M. (2019). Linking mitochondrial dynamics, cristae remodeling and supercomplex formation: How mitochondrial structure can regulate bioenergetics. Mitochondrion, 49, 259–268. 10.1016/j.mito.2019.06.003

Baraibar, A. M., Colomer, T., Moreno-García, A., Bernal-Chico, A., Sánchez-Martín, E., Utrilla, C.,…Mato, S. (2024). Autoimmune inflammation triggers aberrant astrocytic calcium signaling to impair synaptic plasticity. Brain Behav Immun, 121, 192–210. 10.1016/j.bbi.2024.07.010

Barros, L. F., Fernández-Moncada, I., Marsicano, G., Ruminot, I., Saab, A. S., & Weber, B. (2025). Scale-spanning crosstalk between metabolism and information processing. Cell Metab, 37(12), 2303–2310. 10.1016/j.cmet.2025.10.012

Barros, L. F., Ruminot, I., Sotelo-Hitschfeld, T., Lerchundi, R., & Fernández-Moncada, I. (2023). Metabolic recruitment in brain tissue. Annu Rev Physiol, 85, 115–135. 10.1146/annurev-physiol-021422-091035

Beard, E., Lengacher, S., Dias, S., Magistretti, P. J., & Finsterwald, C. (2021). Astrocytes as Key Regulators of Brain Energy Metabolism: New Therapeutic Perspectives. Front Physiol, 12, 825816. 10.3389/fphys.2021.825816

Bittner, C. X., Loaiza, A., Ruminot, I., Larenas, V., Sotelo-Hitschfeld, T., Gutiérrez, R.,…Barros, L. F. (2010). High resolution measurement of the glycolytic rate. Front Neuroenergetics, 2. 10.3389/fnene.2010.00026

Bittner, C. X., Valdebenito, R., Ruminot, I., Loaiza, A., Larenas, V., Sotelo-Hitschfeld, T.,…Barros, L. F. (2011). Fast and reversible stimulation of astrocytic glycolysis by K+ and a delayed and persistent effect of glutamate. J Neurosci, 31(12), 4709–4713. 10.1523/JNEUROSCI.5311-10.2011

Bittner, S., Pape, K., Klotz, L., & Zipp, F. (2023). Implications of immunometabolism for smouldering MS pathology and therapy. Nat Rev Neurol, 19(8), 477–488. 10.1038/s41582-023-00839-6

Bolaños, J. P., & Magistretti, P. J. (2025). The neuron-astrocyte metabolic unit as a cornerstone of brain energy metabolism in health and disease. Nat Metab, 7(12), 2414–2423. 10.1038/s42255-025-01404-9

Bonvento, G., & Bolaños, J. P. (2021). Astrocyte-neuron metabolic cooperation shapes brain activity. Cell Metab, 33(8), 1546–1564. 10.1016/j.cmet.2021.07.006

Bélanger, M., Allaman, I., & Magistretti, P. J. (2011). Brain energy metabolism: focus on astrocyte-neuron metabolic cooperation. Cell Metab, 14(6), 724–738. 10.1016/j.cmet.2011.08.016

Cabral-Costa, J. V., Vicente-Gutiérrez, C., Agulla, J., Lapresa, R., Elrod, J. W., Almeida, Á.,…Kowaltowski, A. J. (2022). Mitochondrial sodium/calcium exchanger NCLX regulates glycolysis in astrocytes, impacting on cognitive performance. J Neurochem. 10.1111/jnc.15745

Cai, P., Li, W., Xu, Y., & Wang, H. (2024). Drp1 and neuroinflammation: Deciphering the interplay between mitochondrial dynamics imbalance and inflammation in neurodegenerative diseases. Neurobiol Dis, 198, 106561. 10.1016/j.nbd.2024.106561

Chao, C. C., Gutiérrez-Vázquez, C., Rothhammer, V., Mayo, L., Wheeler, M. A., Tjon, E. C.,…Quintana, F. J. (2019). Metabolic Control of Astrocyte Pathogenic Activity via cPLA2-MAVS. Cell, 179(7), 1483–1498.e1422. 10.1016/j.cell.2019.11.016

Colomer, T., Bernal-Chico, A., Sánchez-Martín, E., Moreno-García, A., Baraibar, A. M., Uribe-Irusta, A.,…Mato, S. (2026). Astrocyte CB_1_ receptors drive blood-brain barrier disruption in central nervous system inflammatory disease. J Neuroinflammation, 23(1), 73. 10.1186/s12974-026-03708-3

Cunnane, S. C., Trushina, E., Morland, C., Prigione, A., Casadesus, G., Andrews, Z. B.,…Millan, M. J. (2020). Brain energy rescue: an emerging therapeutic concept for neurodegenerative disorders of ageing. Nat Rev Drug Discov, 19(9), 609–633. 10.1038/s41573-020-0072-x

das Neves, S. P., Sousa, J. C., Sousa, N., Cerqueira, J. J., & Marques, F. (2021). Altered astrocytic function in experimental neuroinflammation and multiple sclerosis. Glia, 69(6), 1341–1368. 10.1002/glia.23940

Doni, D., Rigoni, G., Palumbo, E., Baschiera, E., Peruzzo, R., De Rosa, E.,…Costantini, P. (2021). The displacement of frataxin from the mitochondrial cristae correlates with abnormal respiratory supercomplexes formation and bioenergetic defects in cells of Friedreich ataxia patients. FASEB J, 35(3), e21362. 10.1096/fj.202000524RR

Durinck, S., Spellman, P. T., Birney, E., & Huber, W. (2009). Mapping identifiers for the integration of genomic datasets with the R/Bioconductor package biomaRt. Nat Protoc, 4(8), 1184–1191. 10.1038/nprot.2009.97

Dutta, R., McDonough, J., Yin, X., Peterson, J., Chang, A., Torres, T.,…Trapp, B. D. (2006). Mitochondrial dysfunction as a cause of axonal degeneration in multiple sclerosis patients. Ann Neurol, 59(3), 478–489. 10.1002/ana.20736

Eraso-Pichot, A., Pouvreau, S., Olivera-Pinto, A., Gomez-Sotres, P., Skupio, U., & Marsicano, G. (2023). Endocannabinoid signaling in astrocytes. Glia, 71(1), 44–59. 10.1002/glia.24246

Fernández-Moncada, I., Lavanco, G., Fundazuri, U. B., Bollmohr, N., Mountadem, S., Dalla Tor, T.,…Marsicano, G. (2024). A lactate-dependent shift of glycolysis mediates synaptic and cognitive processes in male mice. Nat Commun, 15(1), 6842. 10.1038/s41467-024-51008-2

Gavillet, M., Allaman, I., & Magistretti, P. J. (2008). Modulation of astrocytic metabolic phenotype by proinflammatory cytokines. Glia, 56(9), 975–989. 10.1002/glia.20671

Ghirotto, B., Oliveira, D. F., Cipelli, M., Basso, P. J., de Lima, J., Breda, C. N. S.,…Câmara, N. O. S. (2022). MS-driven metabolic alterations are recapitulated in iPSC-derived astrocytes. Ann Neurol, 91(5), 652–669. 10.1002/ana.26336

Gutiérrez-Rodríguez, A., Bonilla-Del Río, I., Puente, N., Gómez-Urquijo, S. M., Fontaine, C. J., Egaña-Huguet, J.,…Grandes, P. (2018). Localization of the cannabinoid type-1 receptor in subcellular astrocyte compartments of mutant mouse hippocampus. Glia, 66(7), 1417–1431. 10.1002/glia.23314

Gómez-Sotres, P., Skupio, U., Dalla Tor, T., Julio-Kalajzic, F., Cannich, A., Gisquet, D.,…Marsicano, G. (2024). Olfactory bulb astrocytes link social transmission of stress to cognitive adaptation in male mice. Nat Commun, 15(1), 7103. 10.1038/s41467-024-51416-4

Göbel, J., Motori, E., & Bergami, M. (2018). Spatiotemporal control of mitochondrial network dynamics in astroglial cells. Biochem Biophys Res Commun, 500(1), 17–25. 10.1016/j.bbrc.2017.06.191

Hara, T., Amagai, R., Sakakibara, R., & Okado-Matsumoto, A. (2024). Supercomplex formation of mitochondrial respiratory chain complexes in leukocytes from patients with neurodegenerative diseases. J Biochem, 175(3), 289–298. 10.1093/jb/mvad100

Hasel, P., Dando, O., Jiwaji, Z., Baxter, P., Todd, A. C., Heron, S.,…Hardingham, G. E. (2017). Neurons and neuronal activity control gene expression in astrocytes to regulate their development and metabolism. Nat Commun, 8, 15132. 10.1038/ncomms15132

Heo, J. Y., Park, J. H., Kim, S. J., Seo, K. S., Han, J. S., Lee, S. H.,…Kweon, G. R. (2012). DJ-1 null dopaminergic neuronal cells exhibit defects in mitochondrial function and structure: involvement of mitochondrial complex I assembly. PLoS One, 7(3), e32629. 10.1371/journal.pone.0032629

Hertz, L., Peng, L., & Dienel, G. A. (2007). Energy metabolism in astrocytes: high rate of oxidative metabolism and spatiotemporal dependence on glycolysis/glycogenolysis. J Cereb Blood Flow Metab, 27(2), 219–249. 10.1038/sj.jcbfm.9600343

Horvat, A., Muhič, M., Smolič, T., Begić, E., Zorec, R., Kreft, M., & Vardjan, N. (2021). Ca^2+^ as the prime trigger of aerobic glycolysis in astrocytes. Cell Calcium, 95, 102368. 10.1016/j.ceca.2021.102368

Jimenez-Blasco, D., Busquets-Garcia, A., Hebert-Chatelain, E., Serrat, R., Vicente-Gutierrez, C., Ioannidou, C.,…Marsicano, G. (2020). Glucose metabolism links astroglial mitochondria to cannabinoid effects. Nature, 583(7817), 603–608. 10.1038/s41586-020-2470-y

Lange, S. C., Bak, L. K., Waagepetersen, H. S., Schousboe, A., & Norenberg, M. D. (2012). Primary cultures of astrocytes: their value in understanding astrocytes in health and disease. Neurochem Res, 37(11), 2569–2588. 10.1007/s11064-012-0868-0

Lapuente-Brun, E., Moreno-Loshuertos, R., Acín-Pérez, R., Latorre-Pellicer, A., Colás, C., Balsa, E.,…Enríquez, J. A. (2013). Supercomplex assembly determines electron flux in the mitochondrial electron transport chain. Science, 340(6140), 1567–1570. 10.1126/science.1230381

Lassmann, H. (2018). Multiple sclerosis pathology. Cold Spring Harb Perspect Med, 8(3). 10.1101/cshperspect.a028936

Lerchundi, R., Fernández-Moncada, I., Contreras-Baeza, Y., Sotelo-Hitschfeld, T., Mächler, P., Wyss, M. T.,…Barros, L. F. (2015). NH4(+) triggers the release of astrocytic lactate via mitochondrial pyruvate shunting. Proc Natl Acad Sci U S A, 112(35), 11090–11095. 10.1073/pnas.1508259112

Liberzon, A., Birger, C., Thorvaldsdóttir, H., Ghandi, M., Mesirov, J. P., & Tamayo, P. (2015). The Molecular Signatures Database (MSigDB) hallmark gene set collection. Cell Syst, 1(6), 417–425. 10.1016/j.cels.2015.12.004

Licht-Mayer, S., Campbell, G. R., Canizares, M., Mehta, A. R., Gane, A. B., McGill, K.,…Mahad, D. J. (2020). Enhanced axonal response of mitochondria to demyelination offers neuroprotection: implications for multiple sclerosis. Acta Neuropathol, 140(2), 143–167. 10.1007/s00401-020-02179-x

Liddelow, S. A., Guttenplan, K. A., Clarke, L. E., Bennett, F. C., Bohlen, C. J., Schirmer, L.,…Barres, B. A. (2017). Neurotoxic reactive astrocytes are induced by activated microglia. Nature, 541(7638), 481–487. 10.1038/nature21029

Linnerbauer, M., Wheeler, M. A., & Quintana, F. J. (2020). Astrocyte Crosstalk in CNS Inflammation. Neuron, 108(4), 608–622. 10.1016/j.neuron.2020.08.012

Liu, J. J., Mizuno, H., Than-Trong, E., Mächler, P., Kubotera, H., Barbay, T.,…Nasu, Y. (2026). Calcium-independent biosensors for multiplexed *in vivo* imaging of extracellular lactate dynamics. bioRxiv. 10.64898/2026.10.02.756141

Lopez-Fabuel, I., Le Douce, J., Logan, A., James, A. M., Bonvento, G., Murphy, M. P.,…Bolaños, J. P. (2016). Complex I assembly into supercomplexes determines differential mitochondrial ROS production in neurons and astrocytes. Proc Natl Acad Sci U S A, 113(46), 13063–13068. 10.1073/pnas.1613701113

Lopez-Fabuel, I., Martin-Martin, L., Resch-Beusher, M., Azkona, G., Sanchez-Pernaute, R., & Bolaños, J. P. (2017). Mitochondrial respiratory chain disorganization in Parkinson’s disease-relevant PINK1 and DJ1 mutants. Neurochem Int, 109, 101–105. 10.1016/j.neuint.2017.03.023

Lovatt, D., Sonnewald, U., Waagepetersen, H. S., Schousboe, A., He, W., Lin, J. H.,…Nedergaard, M. (2007). The transcriptome and metabolic gene signature of protoplasmic astrocytes in the adult murine cortex. J Neurosci, 27(45), 12255–12266. 10.1523/jneurosci.3404-07.2007

Love, M. I., Huber, W., & Anders, S. (2014). Moderated estimation of fold change and dispersion for RNA-seq data with DESeq2. Genome Biol, 15(12), 550. 10.1186/s13059-014-0550-8

Luo, F., Herrup, K., Qi, X., & Yang, Y. (2017). Inhibition of Drp1 hyper-activation is protective in animal models of experimental multiple sclerosis. Exp Neurol, 292, 21–34. 10.1016/j.expneurol.2017.02.015

Madrer, N., Perera, N. D., Uccelli, N. A., Abbondanza, A., Andersen, J. V., Carsana, E. V.,…Fernández-Moncada, I. (2025). Neural metabolic networks: key elements of healthy brain function. J Neurochem, 169(6), e70084.

Magistretti, P. J., & Allaman, I. (2018). Lactate in the brain: from metabolic end-product to signalling molecule. Nat Rev Neurosci, 19(4), 235–249. 10.1038/nrn.2018.19

Mahad, D. H., Trapp, B. D., & Lassmann, H. (2015). Pathological mechanisms in progressive multiple sclerosis. Lancet Neurol, 14(2), 183–193. 10.1016/S1474-4422(14)70256-X

Mamczur, P., Borsuk, B., Paszko, J., Sas, Z., Mozrzymas, J., Wiśniewski, J. R.,…Rakus, D. (2015). Astrocyte-neuron crosstalk regulates the expression and subcellular localization of carbohydrate metabolism enzymes. Glia, 63(2), 328–340. 10.1002/glia.22753

Marsicano, G., Wotjak, C. T., Azad, S. C., Bisogno, T., Rammes, G., Cascio, M. G.,…Lutz, B. (2002). The endogenous cannabinoid system controls extinction of aversive memories. Nature, 418(6897), 530–534. 10.1038/nature00839

McComish, S. F., O’Sullivan, J., Copas, A. M. M., Imiolek, M., Boyle, N. T., Crompton, L. A.,…Caldwell, M. A. (2024). Reactive astrocytes generated from human iPSC are pro-inflammatory and display altered metabolism. Exp Neurol, 382, 114979. 10.1016/j.expneurol.2024.114979

Meyer, T., Shimon, D., Youssef, S., Yankovitz, G., Tessler, A., Chernobylsky, T.,…Mayo, L. (2022). NAD^+^ metabolism drives astrocyte proinflammatory reprogramming in central nervous system autoimmunity. Proc Natl Acad Sci U S A, 119(35), e2211310119. 10.1073/pnas.2211310119

Moreno-García, A., Serrat, R., Julio-Kalajzic, F., Bernal-Chico, A., Baraibar, A. M., Matute, C.,…Mato, S. (2025). *In vivo* assessment of cortical astrocyte network dysfunction during autoimmune demyelination: correlation with disease severity. J Neurochem, 169(2), e16305. 10.1111/jnc.16305

Moreno-García, Á., Bernal-Chico, A., Colomer, T., Rodríguez-Antigüedad, A., Matute, C., & Mato, S. (2020). Gene expression analysis of astrocyte and microglia endocannabinoid signaling during autoimmune demyelination. Biomolecules, 10(9), 1228. 10.3390/biom10091228

Motori, E., Puyal, J., Toni, N., Ghanem, A., Angeloni, C., Malaguti, M.,…Bergami, M. (2013). Inflammation-induced alteration of astrocyte mitochondrial dynamics requires autophagy for mitochondrial network maintenance. Cell Metab, 18(6), 844–859. 10.1016/j.cmet.2013.11.005

Nasu, Y., Aggarwal, A., Le, G. N. T., Vo, C. T., Kambe, Y., Wang, X.,…Campbell, R. E. (2023). Lactate biosensors for spectrally and spatially multiplexed fluorescence imaging. Nat Commun, 14(1), 6598. 10.1038/s41467-023-42230-5

Ngoc, K. H., Jeon, Y., Ko, J., & Um, J. W. (2025). Multifarious astrocyte-neuron dialog in shaping neural circuit architecture. Trends Cell Biol, 35(1), 74–87. 10.1016/j.tcb.2024.05.002

Nijland, P. G., Michailidou, I., Witte, M. E., Mizee, M. R., van der Pol, S. M., van Het Hof, B.,…van Horssen, J. (2014). Cellular distribution of glucose and monocarboxylate transporters in human brain white matter and multiple sclerosis lesions. Glia, 62(7), 1125–1141. 10.1002/glia.22667

Nijland, P. G., Molenaar, R. J., van der Pol, S. M., van der Valk, P., van Noorden, C. J., de Vries, H. E., & van Horssen, J. (2015). Differential expression of glucose-metabolizing enzymes in multiple sclerosis lesions. Acta Neuropathol Commun, 3, 79. 10.1186/s40478-015-0261-8

Pamies, D., Sartori, C., Schvartz, D., González-Ruiz, V., Pellerin, L., Nunes, C.,…Zurich, M. G. (2021). Neuroinflammatory response to TNFα and IL1β cytokines is accompanied by an increase in glycolysis in human astrocytes in vitro. Int J Mol Sci, 22(8). 10.3390/ijms22084065

Ponath, G., Park, C., & Pitt, D. (2018). The Role of Astrocytes in Multiple Sclerosis. Front Immunol, 9, 217. 10.3389/fimmu.2018.00217

Potter, L. E., Paylor, J. W., Suh, J. S., Tenorio, G., Caliaperumal, J., Colbourne, F.,…Kerr, B. J. (2016). Altered excitatory-inhibitory balance within somatosensory cortex is associated with enhanced plasticity and pain sensitivity in a mouse model of multiple sclerosis. J Neuroinflammation, 13(1), 142. 10.1186/s12974-016-0609-4

Robb, J. L., Hammad, N. A., Weightman Potter, P. G., Chilton, J. K., Beall, C., & Ellacott, K. L. J. (2020). The metabolic response to inflammation in astrocytes is regulated by nuclear factor-kappa B signaling. Glia, 68(11), 2246–2263. 10.1002/glia.23835

Robin, L. M., Oliveira da Cruz, J. F., Langlais, V. C., Martin-Fernandez, M., Metna-Laurent, M., Busquets-Garcia, A.,…Marsicano, G. (2018). Astroglial CB. Neuron, 98(5), 935–944.e935. 10.1016/j.neuron.2018.04.034

Robinson, J. L., Kocabaş, P., Wang, H., Cholley, P. E., Cook, D., Nilsson, A.,…Nielsen, J. (2020). An atlas of human metabolism. Sci Signal, 13(624). 10.1126/scisignal.aaz1482

Rothhammer, V., Mascanfroni, I. D., Bunse, L., Takenaka, M. C., Kenison, J. E., Mayo, L.,…Quintana, F. J. (2016). Type I interferons and microbial metabolites of tryptophan modulate astrocyte activity and central nervous system inflammation via the aryl hydrocarbon receptor. Nat Med, 22(6), 586–597. 10.1038/nm.4106

Russ, K., Teku, G., Bousset, L., Redeker, V., Piel, S., Savchenko, E.,…Roybon, L. (2021). TNF-α and α-synuclein fibrils differently regulate human astrocyte immune reactivity and impair mitochondrial respiration. Cell Rep, 34(12), 108895. 10.1016/j.celrep.2021.108895

San Martín, A., Ceballo, S., Ruminot, I., Lerchundi, R., Frommer, W. B., & Barros, L. F. (2013). A genetically encoded FRET lactate sensor and its use to detect the Warburg effect in single cancer cells. PLoS One, 8(2), e57712. 10.1371/journal.pone.0057712

Schindelin, J., Arganda-Carreras, I., Frise, E., Kaynig, V., Longair, M., Pietzsch, T.,…Cardona, A. (2012). Fiji: an open-source platform for biological-image analysis. Nat Methods, 9(7), 676–682. 10.1038/nmeth.2019

Sotelo-Hitschfeld, T., Niemeyer, M. I., Mächler, P., Ruminot, I., Lerchundi, R., Wyss, M. T.,…Barros, L. F. (2015). Channel-mediated lactate release by K⁺-stimulated astrocytes. J Neurosci, 35(10), 4168–4178. 10.1523/JNEUROSCI.5036-14.2015

Subramanian, A., Tamayo, P., Mootha, V. K., Mukherjee, S., Ebert, B. L., Gillette, M. A.,…Mesirov, J. P. (2005). Gene set enrichment analysis: a knowledge-based approach for interpreting genome-wide expression profiles. Proc Natl Acad Sci U S A, 102(43), 15545–15550. 10.1073/pnas.0506580102

Vicente-Gutierrez, C., Bonora, N., Bobo-Jimenez, V., Jimenez-Blasco, D., Lopez-Fabuel, I., Fernandez, E.,…Bolaños, J. P. (2019). Astrocytic mitochondrial ROS modulate brain metabolism and mouse behaviour. Nat Metab, 1(2), 201–211. 10.1038/s42255-018-0031-6

Véga, C., Pellerin, L., Dantzer, R., & Magistretti, P. J. (2002). Long-term modulation of glucose utilization by IL-1 alpha and TNF-alpha in astrocytes: Na+ pump activity as a potential target via distinct signaling mechanisms. Glia, 39(1), 10–18. 10.1002/glia.10080

Walton, C., King, R., Rechtman, L., Kaye, W., Leray, E., Marrie, R. A.,…Baneke, P. (2020). Rising prevalence of multiple sclerosis worldwide: Insights from the Atlas of MS, third edition. Mult Scler, 26(14), 1816–1821. 10.1177/1352458520970841

Wang, H., Robinson, J. L., Kocabas, P., Gustafsson, J., Anton, M., Cholley, P. E.,…Nielsen, J. (2021). Genome-scale metabolic network reconstruction of model animals as a platform for translational research. Proc Natl Acad Sci U S A, 118(30). 10.1073/pnas.2102344118

Witte, M. E., Bø, L., Rodenburg, R. J., Belien, J. A., Musters, R., Hazes, T.,…van Horssen, J. (2009). Enhanced number and activity of mitochondria in multiple sclerosis lesions. J Pathol, 219(2), 193–204. 10.1002/path.2582

Xiong, J., Ge, X., Pan, D., Zhu, Y., Zhou, Y., Gao, Y.,…Cai, W. (2025). Metabolic reprogramming in astrocytes prevents neuronal death through a UCHL1/PFKFB3/H4K8la positive feedback loop. Cell Death Differ, 32(7), 1214–1230. 10.1038/s41418-025-01467-x

Youle, R. J., & van der Bliek, A. M. (2012). Mitochondrial fission, fusion, and stress. Science, 337(6098), 1062–1065. 10.1126/science.1219855

Zhu, A., Ibrahim, J. G., & Love, M. I. (2019). Heavy-tailed prior distributions for sequence count data: removing the noise and preserving large differences. Bioinformatics, 35(12), 2084–2092. 10.1093/bioinformatics/bty895

